# Nanopore-based genome assembly and the evolutionary genomics of basmati rice

**DOI:** 10.1101/396515

**Authors:** Jae Young Choi, Zoe N. Lye, Simon C. Groen, Xiaoguang Dai, Priyesh Rughani, Sophie Zaaijer, Eoghan D. Harrington, Sissel Juul, Michael D. Purugganan

**Affiliations:** Center for Genomics and Systems Biology, Department of Biology, New York University, New York, New York, USA; Oxford Nanopore Technologies, New York, New York, USA; New York Genome Center, New York, New York, USA; Center for Genomics and Systems Biology, NYU Abu Dhabi Research Institute, New York University Abu Dhabi, Abu Dhabi, United Arab Emirates

**Author notes:** Corresponding authors, (JYC), (MDP).

**Keywords:** *Oryza sativa*, Asian rice, aromatic rice group, domestication, crop evolution, nanopore sequencing, aus, basmati, indica, japonica, admixture, awnless, *de novo* genome assembly

## Abstract

**BACKGROUND:** The *circum-*basmati group of cultivated Asian rice (*Oryza sativa*) contains many iconic varieties and is widespread in the Indian subcontinent. Despite its economic and cultural importance, a high-quality reference genome is currently lacking, and the group’s evolutionary history is not fully resolved. To address these gaps, we used long-read nanopore sequencing and assembled the genomes of two *circum*-basmati rice varieties, Basmati 334 and Dom Sufid.

**RESULTS:** We generated two high-quality, chromosome-level reference genomes that represented the 12 chromosomes of *Oryza*. The assemblies showed a contig N50 of 6.32Mb and 10.53Mb for Basmati 334 and Dom Sufid, respectively. Using our highly contiguous assemblies we characterized structural variations segregating across *circum-*basmati genomes. We discovered repeat expansions not observed in japonica—the rice group most closely related to *circum-* basmati—as well as presence/absence variants of over 20Mb, one of which was a *circum-* basmati-specific deletion of a gene regulating awn length. We further detected strong evidence of admixture between the *circum-*basmati and *circum-*aus groups. This gene flow had its greatest effect on chromosome 10, causing both structural variation and single nucleotide polymorphism to deviate from genome-wide history. Lastly, population genomic analysis of 78 *circum-*basmati varieties showed three major geographically structured genetic groups: (1) Bhutan/Nepal group, (2) India/Bangladesh/Myanmar group, and (3) Iran/Pakistan group.

**CONCLUSION:** Availability of high-quality reference genomes from nanopore sequencing allowed functional and evolutionary genomic analyses, providing genome-wide evidence for gene flow between *circum*-aus and *circum*-basmati, the nature of *circum*-basmati structural variation, and the presence/absence of genes in this important and iconic rice variety group.

## BACKGROUND

*Oryza sativa* or Asian rice is an agriculturally important crop that feeds one-half of the world’s population [1], and supplies 20% of people’s caloric intake (www.fao.org). Historically, *O. sativa* has been classified into two major variety groups, japonica and indica, based on morphometric differences and molecular markers [2, 3]. These variety groups can be considered as subspecies, particularly given the presence of reproductive barriers between them [4]. Archaeobotanical remains suggest japonica rice was domesticated ∼9,000 years ago in the Yangtze Basin of China, while indica rice originated ∼4,000 years ago when domestication alleles were introduced from japonica into either *O. nivara* or a proto-indica in the Indian subcontinent [5]. More recently, two additional variety groups have been recognized that are genetically distinct from japonica and indica: the aus/*circum*-aus and aromatic/*circum*-basmati rices [6–8].

The rich genetic diversity of Asian rice is likely a result from a complex domestication process involving multiple wild progenitor populations and the exchange of important domestication alleles between *O. sativa* variety groups through gene flow [5, 7, 9–17]. Moreover, many agricultural traits within rice are variety group-specific [18–23], suggesting local adaptation to environments or cultural preferences have partially driven the diversification of rice varieties.

Arguably, the *circum*-basmati rice group has been the least studied among the four major variety groups, and it was only recently defined in more detail based on insights from genomic data [7]. Among its members the group boasts the iconic basmati rices (*sensu stricto*) from southern Asia and the sadri rices from Iran [6]. Many, but not all, *circum*-basmati varieties are characterized by distinct and highly desirable fragrance and texture [24]. Nearly all fragrant *circum*-basmati varieties possess a loss-of-function mutation in the *BADH2* gene that has its origins in ancestral japonica haplotypes, suggesting that an introgression between *circum*-basmati and japonica may have led to fragrant basmati rice [21, 25, 26]. Genome-wide polymorphism analysis of a smaller array of *circum*-basmati rice cultivars shows close association with japonica varieties [7, 16, 27], providing evidence that at least part of the genomic make-up of *circum*-basmati rices may indeed be traced back to japonica.

Whole-genome sequences are an important resource for evolutionary geneticists studying plant domestication, as well as breeders aiming to improve crop varieties. Single-molecule sequencing regularly produces sequencing reads in the range of kilobases (kb) [28]. This is particularly helpful for assembling plant genomes, which are often highly repetitive and heterozygous, and commonly underwent at least one round of polyploidization in the past [29–31]. The *Oryza sativa* genome, with a relatively modest size of ∼400 Mb, was the first crop genome sequence assembled [29], and there has been much progress in generating *de novo* genome assemblies for other members of the genus *Oryza*. Currently, there are assemblies for nine wild species (*Leersia perrieri* [outgroup], *O. barthii*, *O. brachyantha*, *O. glumaepatula*, *O. longistaminata*, *O. meridionalis*, *O. nivara*, *O. punctata*, and *O. rufipogon*) and two domesticated species (*O. glaberrima* and *O. sativa*) [32–37].

Within domesticated Asian rice (*O. sativa*), genome assemblies are available for cultivars in most variety groups [32, 33, 38–42]. However, several of these reference assemblies are based on short-read sequencing data and show higher levels of incompleteness compared to assemblies generated from long-read sequences [40, 41]. Nevertheless, these *de novo* genome assemblies have been critical in revealing genomic variation (*e.g.* variations in genome structure and repetitive DNA, and *de novo* species- or population-specific genes) that were otherwise missed from analyzing a single reference genome. Recently, a genome assembly based on short-read sequencing data was generated for basmati rice [42]. Not only were there missing sequences in this assembly, it was also generated from DNA of an elite basmati breeding line. Such modern cultivars are not the best foundations for domestication-related analyses due to higher levels of introgression from other rice populations during modern breeding.

Here, we report the *de novo* sequencing and assembly of the landraces (traditional varieties) Basmati 334 [21, 43, 44] and Dom Sufid [21, 24, 45, 46] using the long-read nanopore sequencing platform of Oxford Nanopore Technologies [47]. Basmati 334 is from Pakistan, evolved in a rainfed lowland environment and is known to be drought tolerant at the seedling and reproductive stages [44]. It also possesses several broad-spectrum bacterial blight resistance alleles [48, 49], making Basmati 334 desirable for breeding resilience into modern basmati cultivars [49, 50]. Dom Sufid is an Iranian sadri cultivar that, like other sadri and basmati (*sensu stricto*) varieties, is among the most expensive varieties currently available in the market [24]. It has desirable characteristics such as aromaticity and grain elongation during cooking, although it is susceptible to disease and abiotic stress [24, 51]. Because of their special characteristics, both Basmati 334 and Dom Sufid are used in elite rice breeding programs to create high yielding and resilient aromatic rice varieties [24, 44–46, 50].

Based on long reads from nanopore sequencing, our genome assemblies have high quality, contiguity, and genic completeness, making them comparable in quality to assemblies associated with key rice reference genomes. We used our *circum*-basmati genome assemblies to characterize genomic variation existing within this important rice variety group, and analyze domestication-related and other evolutionary processes that shaped this variation. Our *circum*-basmati rice genome assemblies will be valuable complements to the available assemblies for other rice cultivars, unlocking important genomic variation for rice crop improvement.

## RESULTS

### Nanopore sequencing of basmati and sadri rice

Using Oxford Nanopore Technologies’ long-read sequencing platform, we sequenced the genomes of the *circum*-basmati landraces Basmati 334 (basmati *sensu stricto*) and Dom Sufid (sadri). We called 1,372,950 reads constituting a total of 29.2 Gb for Basmati 334 and 1,183,159 reads constituting a total of 24.2 Gb for Dom Sufid (Table 1). For both samples the median read length was > 17 kb, the read length N50 was > 33 kb, and the median quality score per read was ∼11.

**Table 1.**
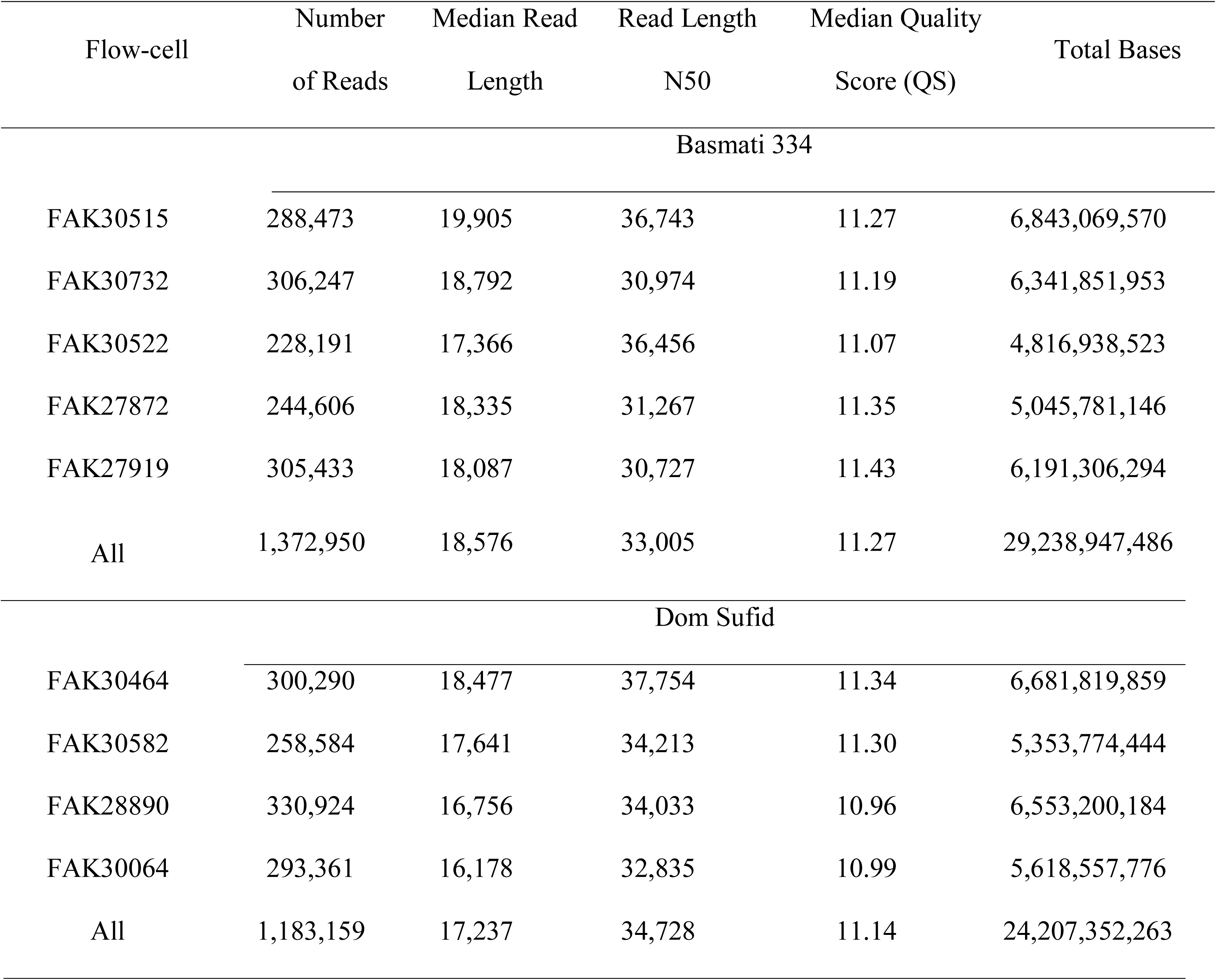
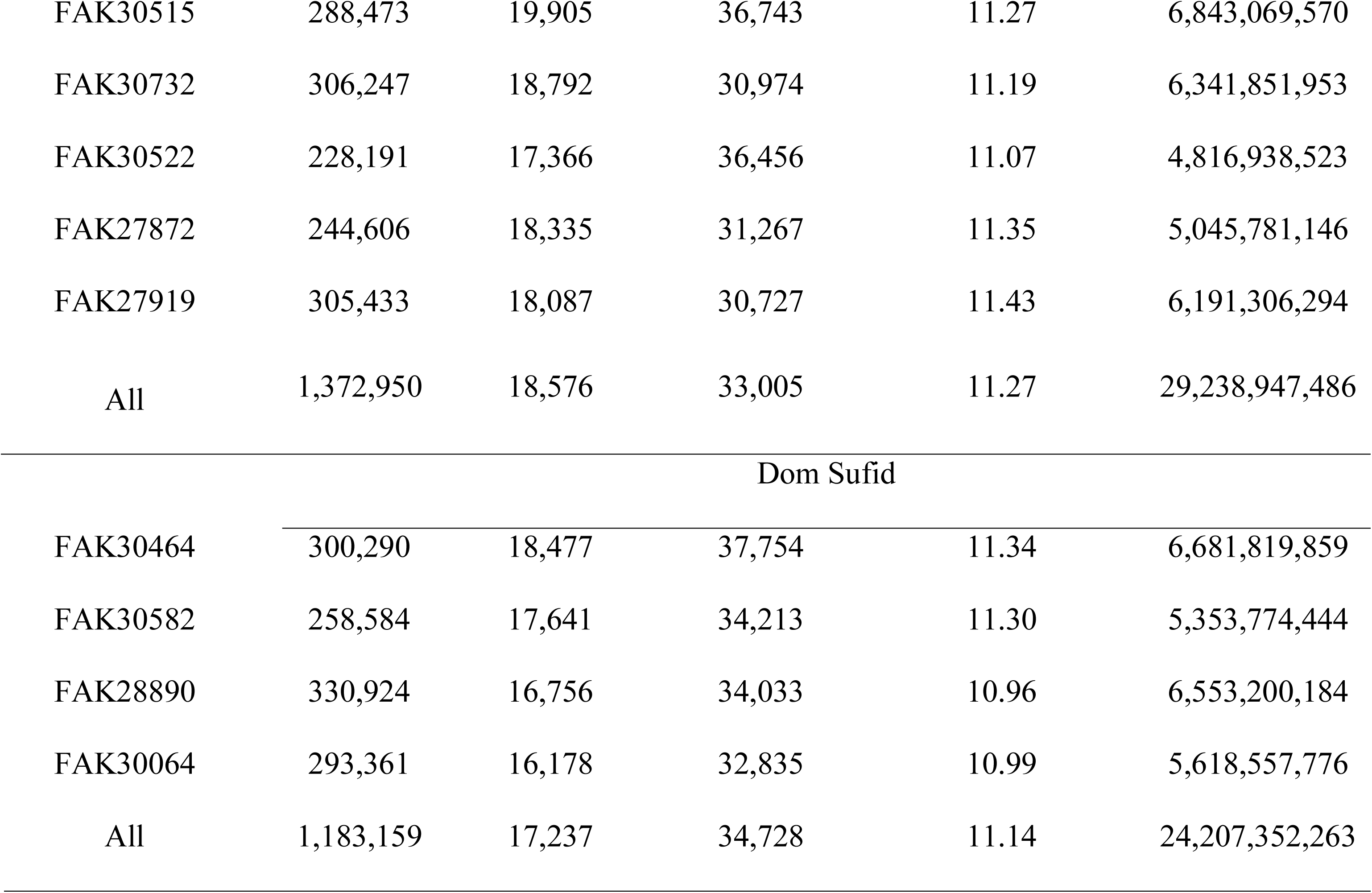
Summary of nanopore sequencing read data.

### *De novo* assembly of the Basmati 334 and Dom Sufid rice genomes

Incorporating only those reads that had a mean quality score of > 8 and read lengths of > 8 kb, we used a total of 1,076,192 reads and 902,040 reads for the Basmati 334 and Dom Sufid genome assemblies, which resulted in a genome coverage of ∼62× and ∼51×, respectively (Table 2). We polished the genome assemblies with both nanopore and short Illumina sequencing reads. The final, polished genome assemblies spanned 386.5 Mb across 188 contigs for Basmati 334, and 383.6 Mb across 116 contigs for Dom Sufid. The genome assemblies had high contiguity, with a contig N50 of 6.32 Mb and 10.53 Mb for Basmati 334 and Dom Sufid, respectively. Our genome assemblies recovered more than 97% of the 1,440 BUSCO [52] embryophyte gene groups, which is comparable to the BUSCO statistics for the japonica Nipponbare [33] (98.4%) and indica R498 reference genomes [41] (98.0%). This is an improvement from the currently available genome assembly of basmati variety GP295-1 [42], which was generated from Illumina short-read sequencing data and has a contig N50 of 44.4 kb with 50,786 assembled contigs.

**Table 2.** Summary of the *circum*-basmati rice genome assemblies.

|  | Basmati 334 | Dom Sufid |
| --- | --- | --- |
| Genome Coverage | 62.5× | 51.4× |
| Number of Contigs | 188 | 116 |
| Total Number of Bases in Contigs | 386,555,741 | 383,636,250 |
| Total Number of Bases Scaffolded | 386,050,525 | 383,245,802 |
| Contig N50 Length | 6.32 Mb | 10.53 Mb |
| Contig L50 | 20 | 13 |
| Total Contigs > 50 kbp | 159 | 104 |
| Maximum Contig Length | 17.04 Mb | 26.82 Mb |
| BUSCO Gene Completion (Assembly) | 97.6% | 97.0% |
| GC Content | 43.6% | 43.7% |
| Repeat Content | 44.4% | 44.2% |
| Number of Annotated Genes | 41,270 | 38,329 |
| BUSCO Gene Completion (Annotation) | 95.4% | 93.6% |

We examined coding sequences of our *circum*-basmati genomes by conducting gene annotation using published rice gene models and the *MAKER* gene annotation pipeline [52, 53]. A total of 41,270 genes were annotated for the Basmati 334 genome, and 38,329 for the Dom Sufid genome. BUSCO gene completion analysis [52] indicated that 95.4% and 93.6% of the 3,278 single copy genes from the liliopsida gene dataset were found in the Basmati 334 and Dom Sufid gene annotations respectively.

### Whole genome comparison to other rice variety group genomes

We aligned our draft genome assemblies to the japonica Nipponbare reference genome sequence [33], which represents one of the highest quality reference genome sequences (Figure 1A). Between the Nipponbare, Basmati 334 and Dom Sufid genomes, high levels of macro-synteny were evident across the japonica chromosomes. Specifically, we observed little large-scale structural variation between Basmati 334 and Dom Sufid contigs and the japonica genome. A noticeable exception was an apparent inversion in the *circum*-basmati genome assemblies at chromosome 6 between positions 12.5 Mb and 18.7 Mb (Nipponbare coordinates), corresponding to the pericentromeric region [54]. Interestingly, the same region showed an inversion between the Nipponbare and indica R498 reference genomes [41], whereas in the *circum*-aus N22 cultivar no inversions are observed (Supplemental Figure 1). While the entire region was inverted in R498, the inversion positions were disjoint in Basmati 334 and Dom Sufid, apparently occurring in multiple regions of the pericentromere. We independently verified the inversions by aligning raw nanopore sequencing reads to the Nipponbare reference genome using the long read-aware aligner *ngmlr* [55], and the structural variation detection program *sniffles* [55]. *Sniffles* detected several inversions, including a large inversion between positions 13.1 and 17.7 Mb and between 18.18 and 18.23 Mb, with several smaller inversions located within the largest inversion (Supplemental Table 1).

**Figure 1.**
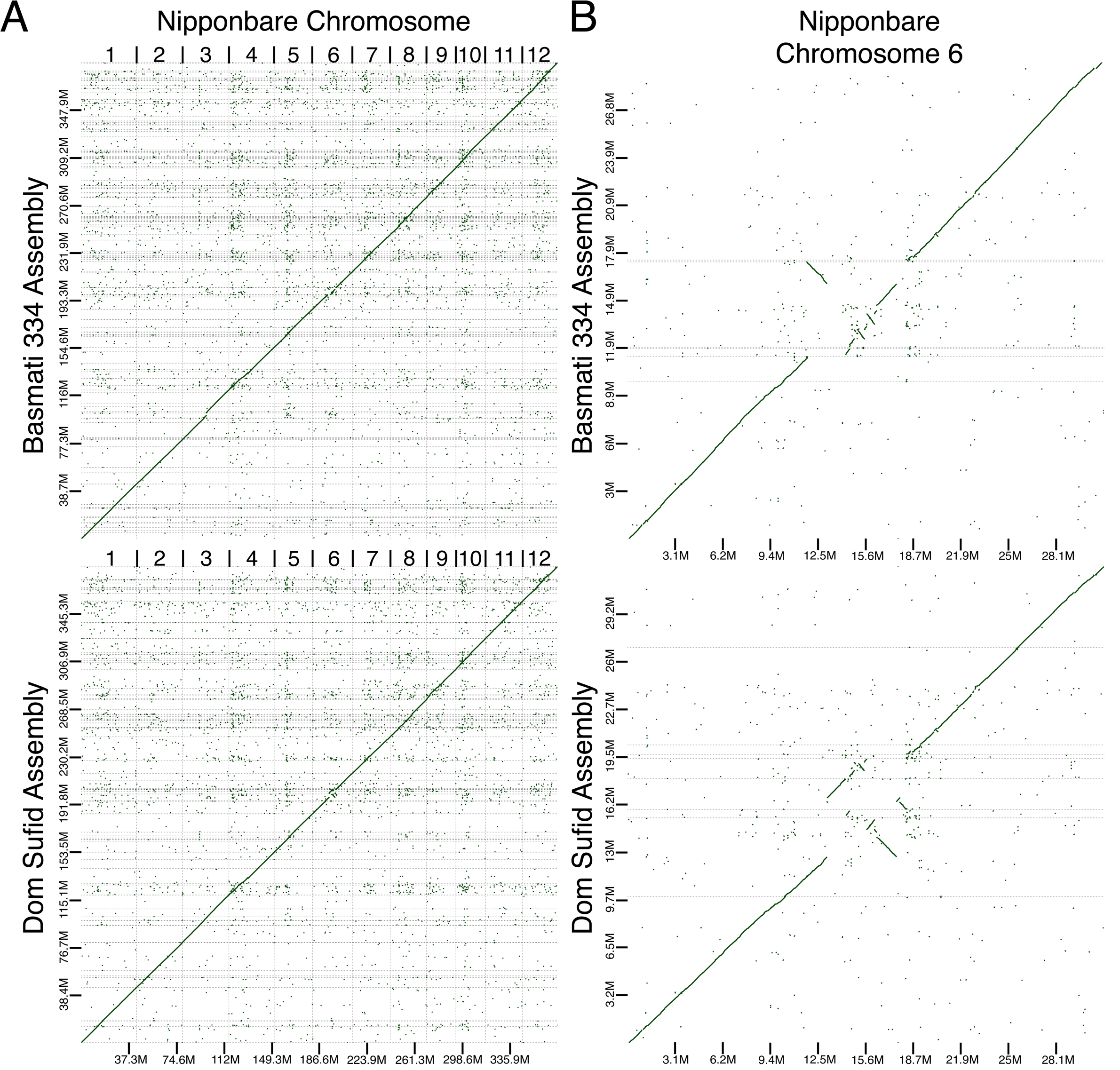
Dot plot comparing the assembly contigs of Basmati 334 and Dom Sufid to (A) all chromosomes of the Nipponbare genome assembly and (B) only chromosome 6 of Nipponbare. Only alignment blocks with greater than 80% overlap in sequence identity are shown.

Because of high macro-synteny with japonica (Figure 1A), we ordered and oriented the contigs of the Basmati 334 and Dom Sufid assemblies using a reference genome-based scaffolding approach [56]. For both Basmati 334 and Dom Sufid, over 99.9% of the assembled genomic contigs were anchored to the Nipponbare reference genome (Table 2). The scaffolded *circum*-basmati chromosomes were similar in size to those in reference genomes for cultivars in other rice variety groups (Nipponbare [33], the *circum*-aus variety N22 [37], and the indica varieties IR8 [37] and R498 [41]) that were sequenced, assembled, and scaffolded to near completion (Table 3).

**Table 3.** Comparison of assembled chromosome sizes for cultivars across variety groups.

| Chromosome | Basmati 334 | Dom Sufid | Nipponbare | N22 | IR8 | R498 |
| --- | --- | --- | --- | --- | --- | --- |
| 1 | 44,411,451 | 44,306,286 | 43,270,923 | 44,711,178 | 44,746,683 | 44,361,539 |
| 2 | 35,924,761 | 36,365,206 | 35,937,250 | 38,372,633 | 37,475,564 | 37,764,328 |
| 3 | 40,305,655 | 38,133,813 | 36,413,819 | 36,762,248 | 39,065,119 | 39,691,490 |
| 4 | 34,905,232 | 34,714,597 | 35,502,694 | 33,558,078 | 35,713,470 | 35,849,732 |
| 5 | 30,669,872 | 31,017,353 | 29,958,434 | 28,792,057 | 31,269,760 | 31,237,231 |
| 6 | 29,982,228 | 32,412,977 | 31,248,787 | 29,772,976 | 32,072,649 | 32,465,040 |
| 7 | 30,410,531 | 29,511,326 | 29,697,621 | 29,936,233 | 30,380,234 | 30,277,827 |
| 8 | 29,921,941 | 29,962,976 | 28,443,022 | 25,527,801 | 30,236,384 | 29,952,003 |
| 9 | 24,050,083 | 23,970,096 | 23,012,720 | 22,277,206 | 24,243,884 | 24,760,661 |
| 10 | 25,596,481 | 24,989,786 | 23,207,287 | 20,972,683 | 25,246,678 | 25,582,588 |
| 11 | 29,979,012 | 29,949,236 | 29,021,106 | 29,032,419 | 32,337,678 | 31,778,392 |
| 12 | 29,893,278 | 27,912,150 | 27,531,856 | 22,563,585 | 25,963,606 | 26,601,357 |
| Total | 386,050,525 | 383,245,802 | 373,245,519 | 362,279,097 | 388,751,709 | 390,322,188 |

Next, we assessed the assembly quality of the *circum*-basmati genomes by contrasting them against available *de novo*-assembled genomes within the Asian rice complex (see Materials and Method for a complete list of genomes). We generated a multi-genome alignment to the Nipponbare genome, which we chose as the reference since its assembly and gene annotation is a product of years of community-based efforts [33, 57, 58]. To infer the quality of the gene regions in each of the genome assemblies, we used the multi-genome alignment to extract the coding DNA sequence of each Nipponbare gene and its orthologous regions from each non-japonica genome. The orthologous genes were counted for missing DNA sequences (“N” sequences) and gaps to estimate the percent of Nipponbare genes covered. For all genomes the majority of Nipponbare genes had a near-zero proportion of sites that were missing in the orthologous non-Nipponbare genes (Supplemental Figure 2). The missing proportions of Nipponbare-orthologous genes within the Basmati 334 and Dom Sufid genomes were comparable to those for genomes that had higher assembly contiguity [37, 40, 41].

Focusing on the previously sequenced basmati GP295-1 genome [42], our newly assembled *circum*-basmati genomes had noticeably lower proportions of missing genes (Supplemental Figure 2). Furthermore, over 96% of base pairs across the Nipponbare genome were alignable against the Basmati 334 (total of 359,557,873 bp [96.33%] of Nipponbare genome) or Dom Sufid (total of 359,819,239 bp [96.40%] of Nipponbare genome) assemblies, while only 194,464,958 bp (52.1%) of the Nipponbare genome were alignable against the GP295-1 assembly.

We then counted the single-nucleotide and insertion/deletion (indel, up to ∼60 bp) differences between the *circum*-basmati and Nipponbare assemblies to assess the overall quality of our newly assembled genomes. To avoid analyzing differences across unconstrained repeat regions, we specifically examined regions where there were 20 exact base-pair matches flanking a site that had a single nucleotide or indel difference between the *circum*-basmati and Nipponbare genomes. In the GP295-1 genome there were 334,500 (0.17%) single-nucleotide differences and 44,609 (0.023%) indels compared to the Nipponbare genome. Our newly assembled genomes had similar proportions of single-nucleotide differences with the Nipponbare genome, where the Basmati 334 genome had 780,735 (0.22%) differences and the Dom Sufid genome had 731,426 (0.20%). For indels the Basmati 334 genome had comparable proportions of differences with 104,282 (0.029%) variants, but the Dom Sufid genome had higher proportions with 222,813 (0.062%) variants. In sum, our draft *circum*-basmati genomes had high contiguity and completeness as evidenced by assembly to the chromosome level, and comparison to the Nipponbare genome. In addition, our genome assemblies were comparable to the Illumina sequence-generated GP295-1 genome for the proportion of genomic differences with the Nipponbare genome, suggesting they had high quality and accuracy as well.

Our *circum*-basmati genome assemblies should also be of sufficiently high quality for detailed gene-level analysis. For instance, a hallmark of many *circum*-basmati rices is aromaticity, and a previous study had determined that Dom Sufid, but not Basmati 334, is a fragrant variety [21]. We examined the two genomes to verify the presence or absence of the mutations associated with fragrance. There are multiple different loss-of-function mutations in the *BADH2* gene that cause rice varieties to be fragrant [21, 25, 26], but the majority of fragrant rices carry a deletion of 8 nucleotides at position chr8:20,382,861-20,382,868 of the Nipponbare genome assembly (version Os-Nipponbare-Reference-IRGSP-1.0). Using the genome alignment, we extracted the *BADH2* sequence region to compare the gene sequence of the non-fragrant Nipponbare to that of Basmati 334 and Dom Sufid. Consistent with previous observations [21], we found that the genome of the non-fragrant Basmati 334 did not carry the deletion and contained the wild-type *BADH2* haplotype observed in Nipponbare. The genome of the fragrant Dom Sufid, on the other hand, carried the 8-bp deletion, as well as the 3 single-nucleotide polymorphisms flanking the deletion. This illustrates that the Basmati 334 and Dom Sufid genomes are accurate enough for gene-level analysis.

### *Circum*-basmati gene analysis

Our annotation identified ∼40,000 coding sequences in the *circum*-basmati assemblies. We examined population frequencies of the annotated gene models across a *circum*-basmati population dataset to filter out mis-annotated gene models or genes at very low frequency in a population. We obtained Illumina sequencing reads from varieties included in the 3K Rice Genome Project [7] and sequenced additional varieties to analyze a total of 78 *circum*-basmati cultivars (see Supplemental Table 2 for a list of varieties). The Illumina sequencing reads were aligned to the *circum*-basmati genomes, and if the average coverage of a genic region was < 0.05× for an individual this gene was called as a deletion in that variety. Since we used a low threshold for calling a deletion, the genome-wide sequencing coverage of a variety did not influence the number of gene deletions detected (Supplemental Figure 3). Results showed that gene deletions were indeed rare across the *circum*-basmati population (Figure 2A), consistent with their probable deleterious nature. We found that 31,565 genes (76.5%) in Basmati 334 and 29,832 genes (77.8%) in the Dom Sufid genomes did not have a deletion across the population (see Supplemental Table 3 for a list of genes).

**Figure 2.**
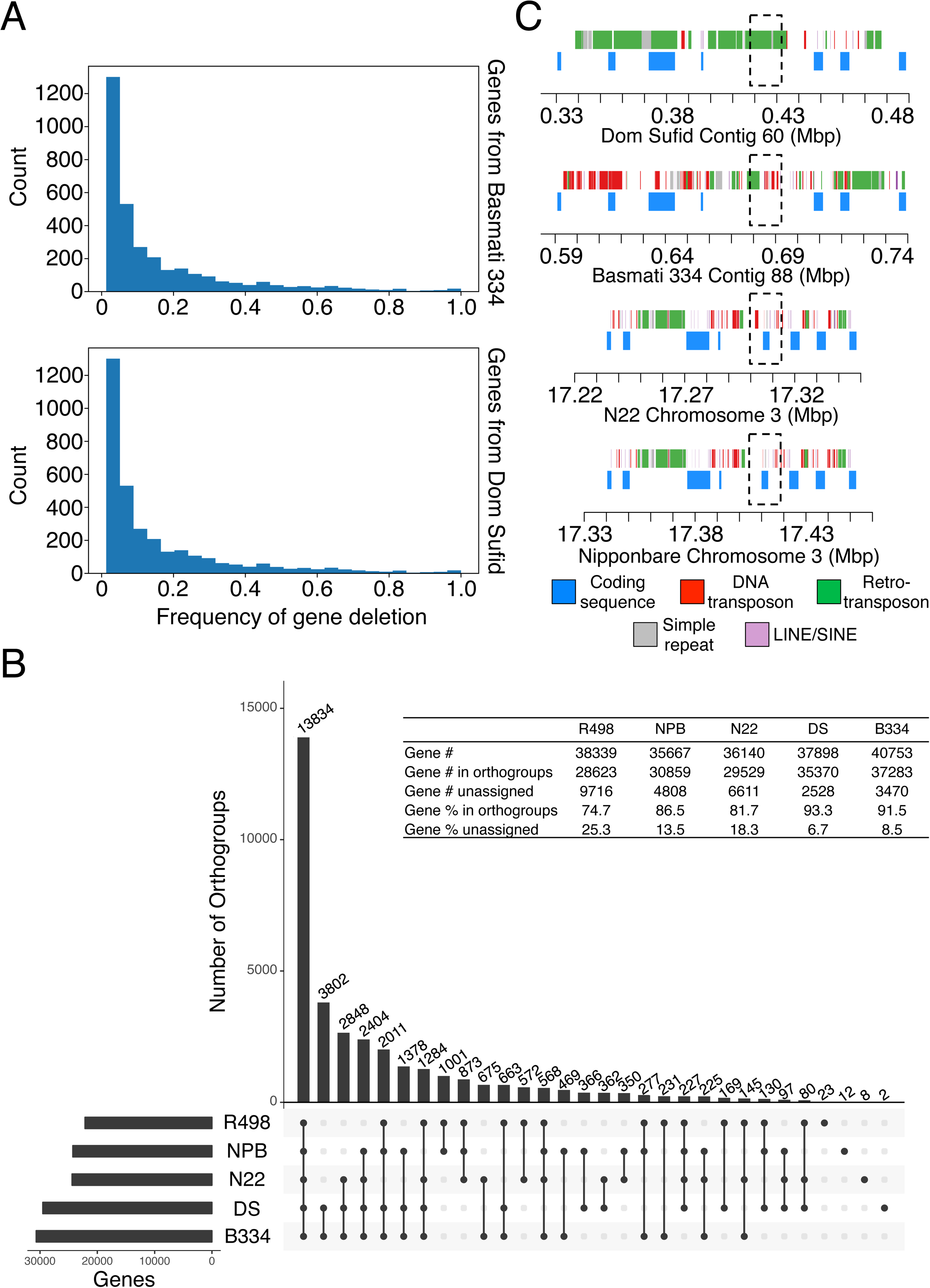
*Circum*-basmati gene sequence evolution. (A) The deletion frequency of genes annotated from the Basmati 334 and Dom Sufid genomes. Frequency was estimated from sequencing data on a population of 78 *circum*-basmati varieties. (B) Groups of orthologous and paralogous genes (*i.e.,* orthogroups) identified in the reference genomes of N22, Nipponbare (NPB), and R498, as well as the *circum*-basmati genome assemblies Basmati 334 (B334) and Dom Sufid (DS) of this study. (C) Visualization of the genomic region orthologous to the Nipponbare gene Os03g0418600 (*Awn3-1*) in the N22, Basmati 334, and Dom Sufid genomes. Regions orthologous to *Awn3-1* are indicated with a dotted box.

There were 517 gene models from Basmati 334 and 431 gene models from Dom Sufid that had a deletion frequency of ≥ 0.3 (see Supplemental Table 4 for a list of genes). These gene ≥ models with high deletion frequencies were not considered further in this analysis. The rest were compared against the *circum*-aus N22, indica R498, and japonica Nipponbare gene models to determine their orthogroup status (Figure 2B; see Supplemental Table 5 for a list of genes and their orthogroup status), which are sets of genes that are orthologs and recent paralogs of each other [59].

The most frequent orthogroup class observed was for groups in which every rice variety group has at least one gene member. There were 13,894 orthogroups within this class, consisting of 17,361 genes from N22, 18,302 genes from Basmati 334, 17,936 genes from Dom Sufid, 17,553 genes from R498, and 18,351 genes from Nipponbare. This orthogroup class likely represents the set of core genes of *O. sativa* [42]. The second-highest orthogroup class observed was for groups with genes that were uniquely found in both *circum*-basmati genomes (3,802 orthogroups). These genes represent those restricted to the *circum*-basmati group.

In comparison to genes in other rice variety groups, the *circum*-basmati genes shared the highest number of orthogroups with *circum*-aus (2,648 orthogroups), followed by japonica (1,378 orthogroups), while sharing the lowest number of orthogroups with indica (663 orthogroups). In fact, genes from indica variety R498 had the lowest number assigned to an orthogroup (Figure 2B inset table), suggesting this genome had more unique genes, *i.e.* without orthologs/paralogs to genes in other rice variety groups.

### Genome-wide presence/absence variation within the *circum*-basmati genomes

Our assembled *circum*-basmati genomes were >10 Mb longer than the Nipponbare genome, but individual chromosomes showed different relative lengths (Table 3) suggesting a considerable number of presence/absence variants (PAVs) between the genomes. We examined the PAVs between the *circum*-basmati and Nipponbare genomes using two different computational packages: (i) *sniffles*, which uses raw nanopore reads aligned to a reference genome to call PAVs, and (ii) *assemblytics* [60], which aligns the genome assemblies to each other and calls PAVs. The results showed that, while the total number of PAVs called by *sniffles* and *assemblytics* were similar, only ∼36% of PAVs had overlapping positions (Table 4). In addition, the combined total size of PAVs was larger for predictions made by *sniffles* compared to those by *assemblytics*. For subsequent analysis we focused on PAVs that were called by both methods.

**Table 4.** Comparison of presence/absence variation called by two different computational packages.

|  | *sniffles* | *assemblytics* | overlap |
| --- | --- | --- | --- |
|  | Basmati 334 |  |  |
| Deletion Counts | 11,989 | 11,247 | 4051 |
| Deleted Basepairs | 43,768,763 | 29,048,238 | 19,328,854 |
| Insertion Counts | 11,447 | 12,161 | 3734 |
| Inserted Basepairs | 19,650,518 | 14,498,550 | 5,783,551 |
|  | Dom Sufid |  |  |
| Deletion Counts | 9901 | 10,115 | 3649 |
| Deleted Basepairs | 36,600,114 | 26,128,143 | 17,274,967 |
| Insertion Counts | 9834 | 11,134 | 3340 |
| Inserted Basepairs | 16,527,995 | 12,902,410 | 5,160,503 |

The distribution of PAV sizes indicated that large PAVs were rare across the *circum*-basmati genomes, while PAVs < 500 bps in size were the most common (Figure 3A). Within smaller-sized PAVs those in the 200-500 bp size range showed a peak in abundance. A closer examination revealed that sequence positions of more than 75% of these 200-500 bp-sized PAVs overlapped with transposable element coordinates in the *circum*-basmati genomes (Supplemental Table 6). A previous study based on short-read Illumina sequencing data reported a similar enrichment of short repetitive elements such as the long terminal repeats (LTRs) of retrotransposons, *Tc1/mariner* elements, and *mPing* elements among PAVs in this size range [61].

**Figure 3.**
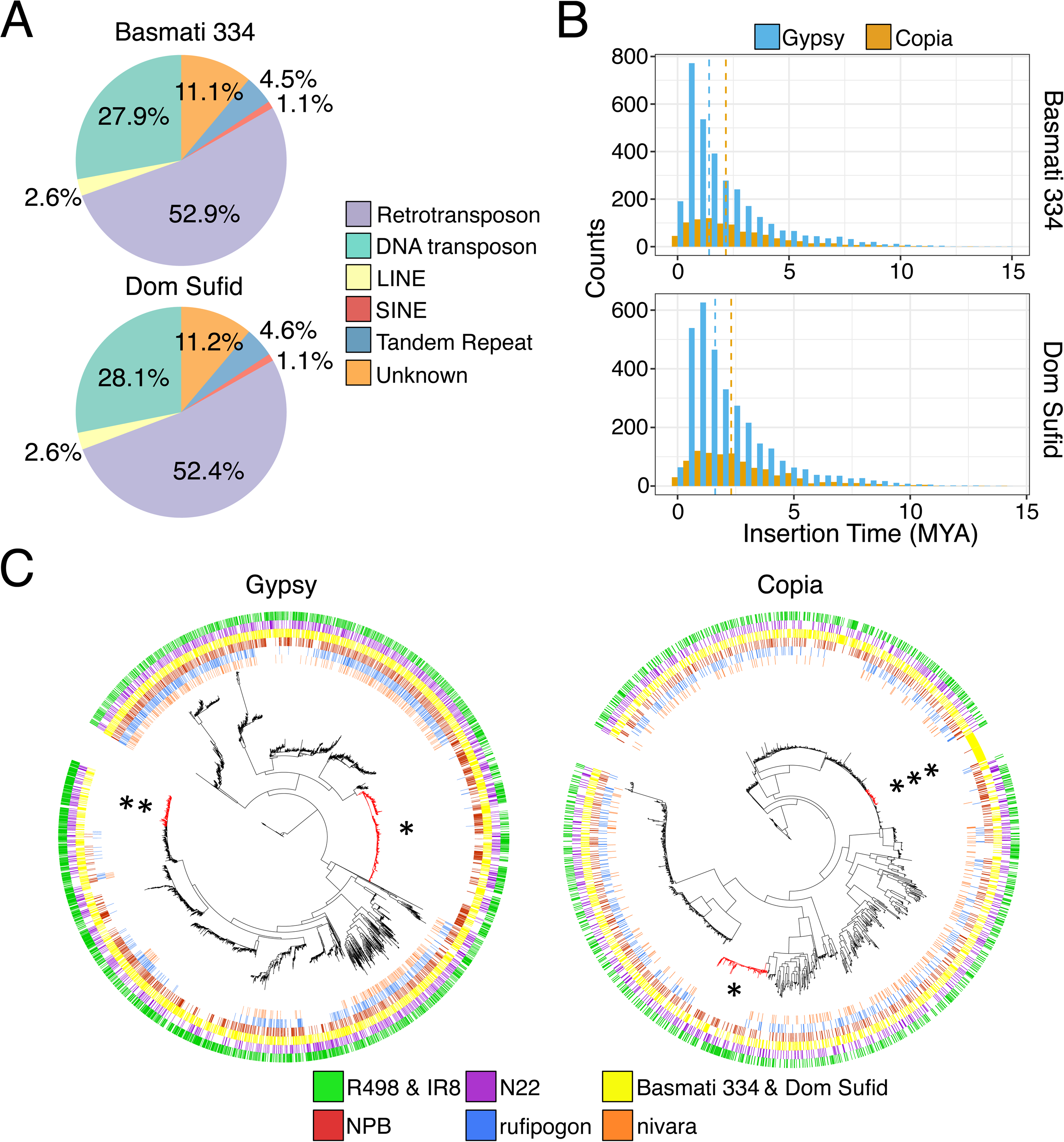
Presence/absence variation across the *circum*-basmati rice genome assemblies. (A) Distribution of presence/absence variant sizes compared to the japonica Nipponbare reference genome. (B) Number of presence/absence variants that are shared between or unique for the *circum*-basmati genomes. (C) Chromosome-wide distribution of presence/absence variation for each *circum*-basmati rice genome, relative to the Nipponbare genome coordinates.

PAVs shorter than 200 bps also overlapped with repetitive sequence positions in the *circum*-basmati genomes, but the relative abundance of each repeat type differed among insertion and deletion variants. Insertions in the Basmati 334 and Dom Sufid genomes had a higher relative abundance of simple sequence repeats (*i.e.* microsatellites) compared to deletions (Supplemental Table 6). These inserted simple sequence repeats were highly enriched for (AT)_n_ dinucleotide repeats, which in Basmati 334 accounted for 66,624 bps out of a total of 72,436 bps (92.0%) of simple sequence repeats, and for Dom Sufid 56,032 bps out of a total of 63,127 bps (88.8%).

Between the Basmati 334 and Dom Sufid genomes, ∼45% of PAVs had overlapping genome coordinates (Figure 3B) suggesting that variety-specific insertion and deletion polymorphisms were common. We plotted PAVs for each of our *circum*-basmati genomes to visualize their distribution (Figure 3C). Chromosome-specific differences in the distribution of PAVs were seen for each *circum*-basmati genome: in Basmati 334, for example, chromosome 1 had the lowest density of PAVs, while in Dom Sufid this was the case for chromosome 2 (Supplemental Figure 5). On the other hand, both genomes showed significantly higher densities of PAVs on chromosome 10 (Tukey’s range test *P* < 0.05). This suggested that, compared to Nipponbare, chromosome 10 was the most differentiated in terms of insertion and deletion variations in both of our *circum*-basmati genomes.

### Evolution of *circum*-basmati rice involved group-specific gene deletions

The proportion of repeat sequences found within the larger-sized PAVs (*i.e.* those > 2 kb) was high, where between 84% and 98% of large PAVs contained transposable element-related sequences (Supplemental Table 6). Regardless, these larger PAVs also involved loss or gain of coding sequences. For instance, gene ontology analysis of domesticated rice gene orthogroups showed enrichment for genes related to electron transporter activity among both *circum*-basmati-specific gene losses and gains (see Supplemental Table 7 for gene ontology results for *circum*-basmati-specific gene losses and Supplemental Table 8 for gene ontology results for *circum*-basmati-specific gene gains).

Many of these genic PAVs could have been important during the rice domestication process [11]. Gene deletions, in particular, are more likely to have a functional consequence than single-nucleotide polymorphisms or short indels and may underlie drastic phenotypic variation. In the context of crop domestication and diversification this could have led to desirable phenotypes in human-created agricultural environments. For instance, several domestication phenotypes in rice are known to be caused by gene deletions [35, 62–66].

There were 873 gene orthogroups for which neither of the *circum*-basmati genomes had a gene member, but for which genomes for all three other rice variety groups (N22, Nipponbare, and R498) had at least one gene member. Among these, there were 545 orthogroups for which N22, Nipponbare, and R498 each had a single-copy gene member, suggesting that the deletion of these genes in both the Basmati 334 and Dom Sufid genomes could have had a major effect in *circum*-basmati. We aligned Illumina sequencing data from our *circum*-basmati population dataset to the japonica Nipponbare genome, and calculated deletion frequencies of Nipponbare genes that belonged to the 545 orthogroups (see Supplemental Table 9 for gene deletion frequencies in the *circum*-basmati population for the Nipponbare genes that are missing in Basmati 334 and Dom Sufid). The vast majority of these Nipponbare genes (509 orthogroups or 93.4%) were entirely absent in the *circum*-basmati population, further indicating that these were *circum*-basmati-specific gene deletions fixed within this variety group.

One of the genes specifically deleted in *circum*-basmati rice varieties was *Awn3-1* (Os03g0418600), which was identified in a previous study as associated with altered awn length in japonica rice [67]. Reduced awn length is an important domestication trait that was selected for ease of harvesting and storing rice seeds [68]. This gene was missing in both *circum*-basmati genomes and no region could be aligned to the Nipponbare *Awn3-1* genic region (Figure 2C). Instead of the *Awn3-1* coding sequence, this genomic region contained an excess of transposable element sequences, suggesting an accumulation of repetitive DNA may have been involved in this gene’s deletion. The flanking arms upstream and downstream of Os03g0418600 were annotated in both *circum*-basmati genomes and were syntenic to the regions in both Nipponbare and N22. These flanking arms, however, were also accumulating transposable element sequences, indicating that this entire genomic region may be degenerating in both *circum*-basmati rice genomes.

### Repetitive DNA and retrotransposon dynamics in the *circum*-basmati genomes

Repetitive DNA makes up more than 44% of the Basmati 334 and Dom Sufid genome assemblies (Table 2). Consistent with genomes of other plant species [69] the repetitive DNA was largely composed of Class I retrotransposons, followed by Class II DNA transposons (Figure 4A). In total, 171.1 Mb were annotated as repetitive for Basmati 334, and 169.5 Mb for Dom Sufid. The amount of repetitive DNA in the *circum*-basmati genomes was higher than in the Nipponbare (160.6 Mb) and N22 genomes (152.1 Mb), but lower than in the indica R498 (175.9 Mb) and IR8 (176.0 Mb) genomes. These differences in the total amount of repetitive DNA were similar to overall genome assembly size differences (Table 3), indicating that variation in repeat DNA accumulation is largely driving genome size differences in rice [70].

**Figure 4.**
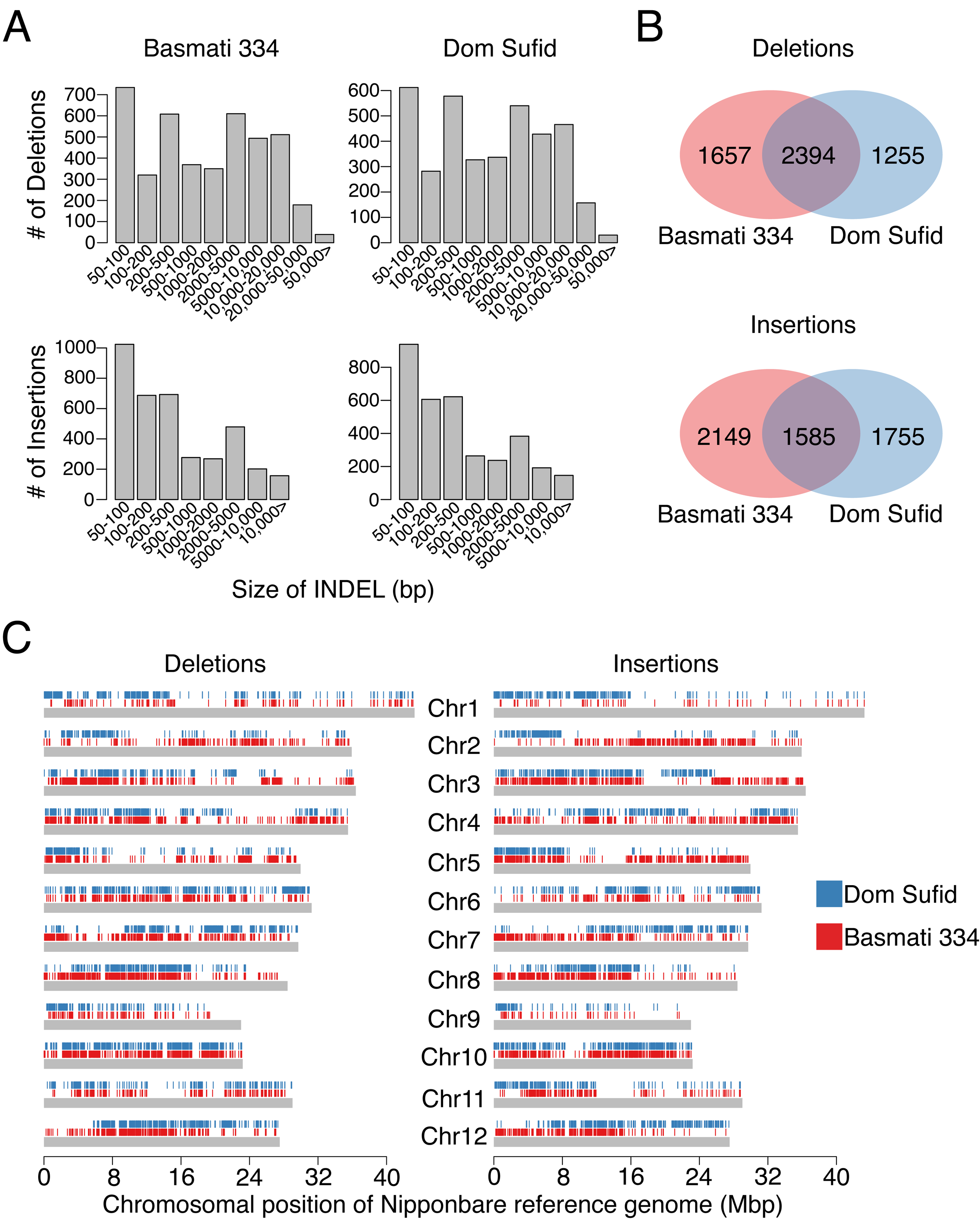
Repetitive DNA landscape of the Basmati 334 and Dom Sufid genomes. (A) Proportion of repetitive DNA content in the *circum*-basmati genomes represented by each repeat family. (B) Distribution of insert times for the *gypsy* and *copia* LTR retrotransposons. (C) Phylogeny of *gypsy* and *copia* LTR retrotransposons based on the *rve* gene. LTR retrotransposons were annotated from the reference genomes of domesticated and wild rices.

We focused our attention on retrotransposons, which made up the majority of the rice repetitive DNA landscape (Figure 4A). Using *LTRharvest* [71, 72], we identified and *de novo*-annotated LTR retrotransposons in the *circum*-basmati genomes. *LTRharvest* annotated 5,170 and 5,150 candidate LTR retrotransposons in Basmati 334 and Dom Sufid, respectively (Supplemental Tables 10 and 11). Of these, 4,180 retrotransposons (80.9% of all candidate LTR retrotransposons) in Basmati 334 and 4,228 (82.1%) in Dom Sufid were classified as LTR retrotransposons by *RepeatMasker*’s *RepeatClassifer* tool (http://www.repeatmasker.org). Most LTR retrotransposons were from the *gypsy* and *copia* superfamilies [73, 74], which made up 77.1% (3,225 *gypsy* elements) and 21.9% (915 *copia* elements) of LTR retrotransposons in the Basmati 334 genome, and 76.4% (3,231 *gypsy* elements) and 22.8% (962 *copia* elements) of LTR retrotransposons in the Dom Sufid genome, respectively. Comparison of LTR retrotransposon content among reference genomes from different rice variety groups (Supplemental Figure 4) revealed that genomes assembled to near completion (*i.e.* Nipponbare, N22, Basmati 334, Dom Sufid, and indica varieties IR8 and R498, as well as MH63 and ZS97 [40]) had higher numbers of annotated retrotransposons than genomes generated from short-read sequencing data (GP295-1, *circum*-aus varieties DJ123 [38] and Kasalath [39], and indica variety IR64 [38]), suggesting genome assemblies from short-read sequencing data may be missing certain repetitive DNA regions.

Due to the proliferation mechanism of LTR transposons, the DNA divergence of an LTR sequence can be used to approximate the insertion time for an LTR retrotransposon [75]. Compared to other rice reference genomes, the insertion times for the Basmati 334 and Dom Sufid LTR retrotransposons were most similar to those observed for elements in the *circum*-aus N22 genome (Supplemental Figure 4). Within our *circum*-basmati assemblies, the *gypsy* superfamily elements had a younger average insertion time (∼2.2 million years ago) than elements of the *copia* superfamily (∼2.7 million years ago; Figure 4B).

Concentrating on *gypsy* and *copia* elements with the *rve* (integrase; Pfam ID: PF00665) gene, we examined the evolutionary dynamics of these LTR retrotransposons by reconstructing their phylogenetic relationships across reference genomes for the four domesticated rice variety groups (N22, Basmati 334, Dom Sufid, R498, IR8, and Nipponbare), and the two wild rice species (*O. nivara* and *O. rufipogon*; Fig 3C). The retrotransposons grouped into distinct phylogenetic clades, which likely reflect repeats belonging to the same family or subfamily [76]. The majority of phylogenetic clades displayed short external and long internal branches, consistent with rapid recent bursts of transposition observed across various rice LTR retrotransposon families [77].

The *gypsy* and *copia* superfamilies each contained a clade in which the majority of elements originated within *O. sativa*, present only among the four domesticated rice variety groups (Figure 4C, single star; see Supplemental Tables 12 and 13 for their genome coordinates). Elements in the *gypsy* superfamily phylogenetic clade had sequence similarity (963 out of the 1,837 retrotransposons) to elements of the *hopi* family [78], while elements in the *copia* superfamily phylogenetic clade had sequence similarity (88 out of the 264) to elements in the *osr4* family [79]. Elements of the *hopi* family are found in high copy number in genomes of domesticated rice varieties [80] and this amplification has happened recently [81].

Several retrotransposon clades were restricted to certain rice variety groups. The *gypsy* superfamily harbored a phylogenetic clade whose elements were only present in genomes of *circum*-aus, *circum*-basmati, and indica varieties (Figure 4C, double star; see Supplemental Table 14 for their genome coordinates), while we observed a clade comprised mostly of *circum*-basmati-specific elements within the *copia* superfamily (Figure 4C, triple star; see Supplemental Table 15 for their genome coordinates). Only a few members of the *gypsy*-like clade had sequence similarity (7 out of 478) to elements of the *rire3* [82] and *rn215* [83] families. Members of both families are known to be present in high copy numbers in genomes of domesticated rice varieties, but their abundance differs between the japonica and indica variety groups [80], suggesting a *rire3*- or *rn215*-like element expansion in the *circum*-aus, *circum*-basmati, and indica genomes. A majority of the *circum*-basmati-specific *copia*-like elements had sequence similarity (109 out of 113) to members of the *houba* family [78], which are found in high copy numbers in certain individuals, but in lower frequency across the rice population [80]. This suggests the *houba* family might have undergone a recent expansion specifically within the *circum*-basmati genomes.

### Phylogenomic analysis on the origins of *circum*-basmati rice

We estimated the phylogenetic relationships within and between variety groups of domesticated Asian rice. Our maximum-likelihood phylogenetic tree, based on four-fold degenerate sites from the Nipponbare coding sequences (Figure 5A), showed that each cultivar was monophyletic with respect to its variety group of origin. In addition, the *circum*-basmati group was sister to japonica rice, while the *circum*-aus group was sister to indica. Consistent with previous observations, the wild rices *O. nivara* and *O. rufipogon* were sister to the *circum*-aus and japonica rices, respectively [14]. While this suggests that each domesticated rice variety group may have had independent wild progenitors of origin, it should be noted that recent hybridization between wild and domesticated rice [84, 85] could lead to similar phylogenetic relationships.

**Figure 5.**
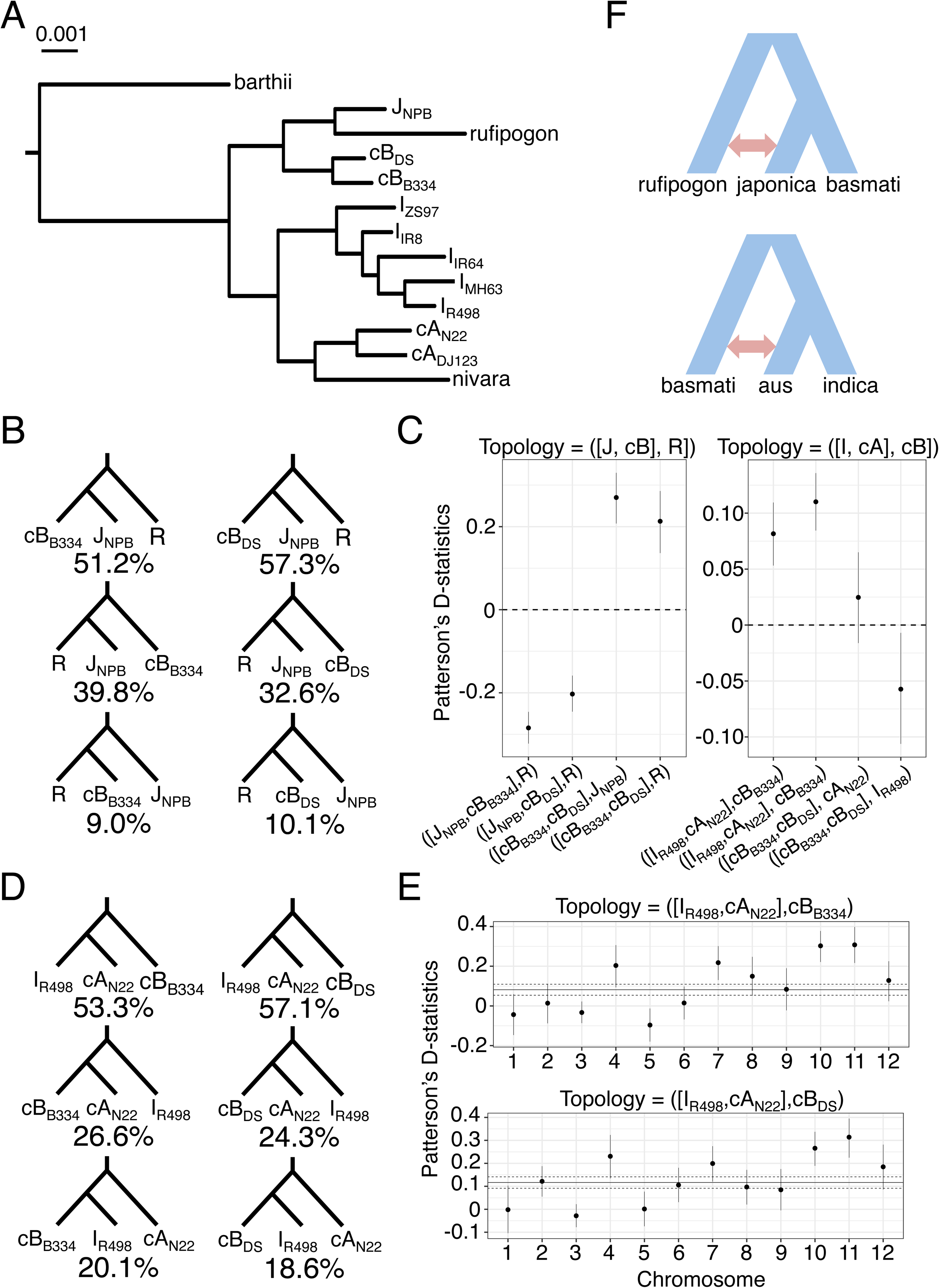
Comparative genomic analysis of *circum*-basmati rice evolution. (A) Maximum-likelihood tree based on four-fold degenerate sites. All nodes had over 95% bootstrap support. (B) Percentage of genes supporting the topology involving japonica (J; Nipponbare, NPB), *circum*-basmati (cB, *circum*-basmati; Basmati 334, B334; Dom Sufid, DS), and *O. rufipogon* (R) after an Approximately Unbiased (AU) test. (C) Results of ABBA-BABA tests. Shown are median Patterson’s D-statistics with 95% confidence intervals determined from a bootstrapping procedure. For each tested topology the outgroup was always *O. barthii*. (D) Percentage of genes supporting the topology involving *circum*-aus (cA; N22), *circum*-basmati, and indica (I; R498) after an Approximately Unbiased (AU) test. (E) Per-chromosome distribution of D-statistics for the trio involving R498, N22, and each *circum*-basmati genome. Genome-wide D-statistics with 95% bootstrap confidence intervals are indicated by the dark and dotted lines. (F) Model of admixture events that occurred within domesticated Asian rice. The direction of admixture has been left ambiguous as the ABBA-BABA test cannot detect the direction of gene flow. The *Oryza sativa* variety groups are labeled as *circum*-aus (cA), *circum*-basmati (cB), indica (I), and japonica (J), and the wild relative is *O. rufipogon* (R).

To further investigate phylogenetic relationships between *circum*-basmati and japonica, we examined phylogenetic topologies of each gene involving the trio Basmati 334, Nipponbare, and *O. rufipogon*. For each gene we tested which of three possible topologies for a rooted three-species tree - *i.e.* [(P1, P2), P3], O, where O is outgroup *O. barthii* and P1, P2, and P3 are Basmati 334 (or Dom Sufid), Nipponbare, and *O. rufipogon*, respectively - were found in highest proportion. For the trio involving Basmati 334, Nipponbare, and *O. rufipogon* there were 7,581 genes (or 32.6%), and for the trio involving Dom Sufid, Nipponbare, and *O. rufipogon* there were 7,690 genes (or 33.1%), that significantly rejected one topology over the other two using an Approximately Unbiased (AU) topology test [86]. In both trios, the majority of those genes supported a topology that grouped *circum*-basmati and Nipponbare as sister to each other (Figure 5B; 3,881 [or 51.2%] and 4,407 [or 57.3%] genes for Basmati 334 and Dom Sufid, respectively). A lower number of genes (3,018 [or 39.8%] and 2,508 [or 32.6%] genes for Basmati 334 and Dom Sufid, respectively) supported the topology that placed Nipponbare and *O. rufipogon* together.

The topology test result suggested that [(japonica, *circum*-basmati), *O. rufipogon*] was the true species topology, while the topology [(japonica, *O. rufipogon*), *circum*-basmati)] represented possible evidence of admixture (although it could also arise from incomplete lineage sorting). To test for introgression, we employed D-statistics from the ABBA-BABA test [87, 88]. The D-statistics for the topology [(japonica, *circum*-basmati), *O. rufipogon*] were significantly negative - Figure 5C left panel; z-score = −14.60 and D ± s.e = −0.28 ± 0.019 for topology [(Nipponbare, Basmati 334), *O. rufipogon*], and z-score = −9.09 and D = −0.20 ± 0.022 for topology [(Nipponbare, Dom Sufid), *O. rufipogon*] - suggesting significant evidence of admixture between japonica and *O. rufipogon*.

Our initial topology test suggested that the trio involving Dom Sufid, Nipponbare, and *O. rufipogon* had a higher proportion of genes supporting the [(*circum*-basmati, japonica), *O. rufipogon*] topology compared to the trio involving Basmati 334, Nipponbare, and *O. rufipogon* (Figure 5B). This suggested within-population variation in the amount of japonica or *O. rufipogon* ancestry across the *circum*-basmati genomes due to differences in gene flow. We conducted ABBA-BABA tests involving the topology [(Basmati 334, Dom Sufid), Nipponbare or *O. rufipogon*] to examine the differences in introgression between the *circum*-basmati and japonica or *O. rufipogon* genomes. The results showed significantly positive D-statistics for the topology [(Basmati 334, Dom Sufid), Nipponbare] (Figure 5C left panel; z-score = 8.42 and D = 0.27 ± 0.032), indicating that Dom Sufid shared more alleles with japonica than Basmati 334 did due to a history of more admixture with japonica. The D-statistics involving the topology [(Basmati 334, Dom Sufid), *O. rufipogon*] were also significantly positive (Figure 5C left panel; z-score = 5.57 and D = 0.21 ± 0.038). While this suggests admixture between Dom Sufid and *O. rufipogon*, it may also be an artifact due to the significant admixture between japonica and *O. rufipogon*.

### Signatures of admixture between *circum*-basmati and *circum*-aus rice genomes

Due to extensive admixture between rice variety group genomes [14] we examined whether the basmati genome was also influenced by gene flow with other divergent rice variety groups (*i.e. circum*-aus or indica rices). A topology test was conducted for a rooted, three-population species tree. For the trio involving Basmati 334, *circum*-aus variety N22, and indica variety R498 there were 7,859 genes (or 35.3%), and for the trio involving Dom Sufid, N22, and R498 there were 8,109 genes (or 37.8%), that significantly rejected one topology over the other two after an AU test. In both trios, more than half of the genes supported the topology grouping *circum*-aus and indica as sisters (Figure 5D). In addition, more genes supported the topology grouping *circum*-aus and *circum*-basmati as sisters than the topology grouping indica and *circum*-basmati as sisters. This suggested that the *circum*-aus variety group might have contributed a larger proportion of genes to *circum*-basmati through gene flow than the indica variety group did.

To test for evidence of admixture, we conducted ABBA-BABA tests involving trios of the *circum*-basmati, N22, and R498 genomes. Results showed significant evidence of gene flow between *circum*-aus and both *circum*-basmati genomes - Figure 5C, right panel; z-score = 5.70 and D = 0.082 ± 0.014 for topology [(R498, N22), Basmati 334]; and z-score = 8.44 and D = 0.11 ± 0.013 for topology [(R498, N22), Dom Sufid]. To test whether there was variability in the *circum*-aus or indica ancestry in each of the *circum*-basmati genomes, we conducted ABBA-BABA tests for the topology [(Basmati 334, Dom Sufid), N22 or R498]. Neither of the ABBA-BABA tests involving the topology [(Basmati 334, Dom Sufid), N22] (Figure 5C, right panel; z-score = 1.20 and D = 0.025 ± 0.021) or the topology [(Basmati 334, Dom Sufid), R498) (Figure 5C, right panel; z-score = −2.24 and D = −0.06 ± 0.026) was significant, suggesting the amount of admixture from *circum*-aus to each of the two *circum*-basmati genomes was similar.

In sum, the phylogenomic analysis indicated that *circum*-basmati and japonica share the most recent common ancestor, while *circum*-aus has admixed with *circum*-basmati during its evolutionary history (Figure 5F). We then examined whether admixture from *circum*-aus had affected each of the *circum*-basmati chromosomes to a similar degree. For both *circum*-basmati genomes most chromosomes had D-statistics that were not different from the genome-wide D-statistics value or from zero (Figure 5E). Exceptions were chromosomes 10 and 11, where the bootstrap D-statistics were significantly higher than the genome-wide estimate.

### Population analysis on the origin of *circum*-basmati rice

Since our analysis was based on single representative genomes from each rice variety group, we compared the results of our phylogenomic analyses to population genomic patterns in an expanded set of rice varieties from different groups. We obtained high coverage (>14×) genomic re-sequencing data (generated with Illumina short-read sequencing) from landrace varieties in the 3K Rice Genome Project [7] and from *circum*-basmati rice landraces we re-sequenced. In total, we analyzed 24 *circum*-aus, 18 *circum*-basmati, and 37 tropical japonica landraces (see Supplemental Table 16 for variety names). The raw Illumina sequencing reads were aligned to the scaffolded Basmati 334 genome and computationally genotyped. A total of 4,594,290 polymorphic sites were called across the three rice variety groups and used for further analysis.

To quantify relationships between *circum*-aus, *circum*-basmati, and japonica, we conducted a topology-weighting analysis [89]. For three populations there are three possible topologies and we conducted localized sliding window analysis to quantify the number of unique sub-trees that supported each tree topology. Consistent with the phylogenomic analysis results, the topology weight was largest for the topology that grouped japonica and *circum*-basmati as sisters (Figure 6A; topology weight = 0.481 with 95% confidence interval [0.479-0.483]). The topology that grouped *circum*-aus and *circum*-basmati together as sisters weighed significantly more (topology weight = 0.318 with 95% confidence interval [0.316-0.320]) than the topology that grouped japonica and *circum*-aus as sisters (topology weight = 0.201 with 95% confidence interval [0.199-0.203). This was consistent with the admixture results from the comparative phylogenomic analysis, which detected evidence of gene flow between *circum*-aus and *circum*-basmati.

**Figure 6.**
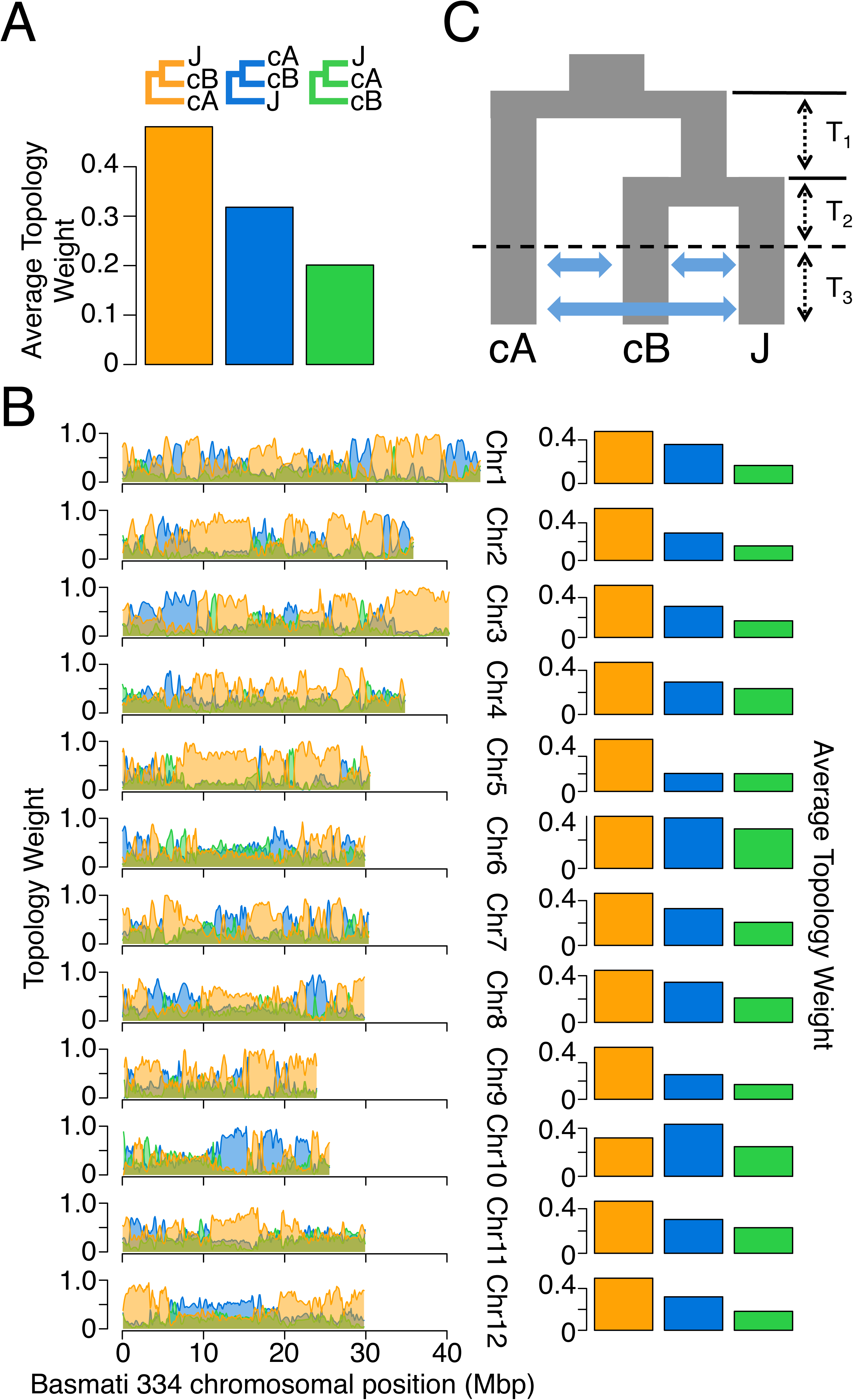
Population relationships among the *circum*-aus (cA), *circum*-basmati (cB), and japonica rices (J). (A) Sum of genome-wide topology weights for a three-population topology involving trios of the *circum*-aus, *circum*-basmati, and japonica rices. Topology weights were estimated across windows with 100 SNPs. (B) Chromosomal distributions of topology weights involving trios of the *circum*-aus, *circum*-basmati, and japonica rices (left), and the sum of the topology weights (right). (C) Best-fitting *δaδi* model for the *circum*-aus, *circum*-basmati, and japonica rices. See Supplemental Table 17 for parameter estimates.

We then examined topology weights for each individual chromosome, since the ABBA-BABA tests using the genome assemblies had detected variation in *circum*-aus ancestry between different chromosomes (Figure 5E). The results showed that for most of the chromosomes the topology [(japonica, *circum*-basmati), *circum*-aus] always weighed more than the remaining two topologies. An exception was observed for chromosome 10 where the topology weight grouping *circum*-aus and *circum*-basmati as sisters was significantly higher (topology weight = 0.433 with 95% confidence interval [0.424-0.442]) than the weight for the genome-wide topology that grouped japonica and *circum*-basmati as sisters (topology weight = 0.320 with 95% confidence interval [0.312-0.328]). This change in predominant topology was still observed when the weights were calculated across wider local windows (Supplemental Figure 6). Another exception could be seen for chromosome 6 where the genome-wide topology [(japonica, *circum*-basmati), *circum*-aus] (topology weight = 0.367 with 95% confidence interval [0.359-0.374) and the admixture topology [*circum*-aus, *circum*-basmati), japonica] (topology weight = 0.355 with 95% confidence interval [0.349-0.362]) had almost equal weights. In larger window sizes the weight of the admixed topology was slightly higher than that of the genome-wide topology (Supplemental Figure 6).

To estimate the evolutionary/domestication scenario that might explain the observed relationships between the *circum*-aus, *circum*-basmati, and japonica groups, we used the diffusion-based approach of the program *δaδi* [90] and fitted specific demographic models to the observed allele frequency spectra for the three rice variety groups. Because all three rice groups have evidence of admixture with each other [7, 9, 14, 16] we examined 13 demographic scenarios involving symmetric, asymmetric, and “no migration” models between variety groups, with and without recent population size changes (Supplemental Figure 7). To minimize the effect of genetic linkage on the demography estimation, polymorphic sites were randomly pruned in 200 kb windows, resulting in 1,918 segregating sites. The best-fitting demographic scenario was one that modeled a period of lineage splitting and isolation, while gene flow only occurred after formation of the three populations and at a later time (Figure 6C; visualizations of the 2D site frequency spectrum and model fit can be seen in Supplemental Figure 8). This best-fitting model was one of the lesser-parameterized models we tested, and the difference in Akaike Information Criterion (ΔAIC) with the model with the second-highest likelihood was 25.46 (see Δ Supplemental Table 17 for parameter estimates and maximum likelihood estimates for each demographic model).

### Genetic structure within the *circum*-basmati group

We used the *circum*-basmati population genomic data for the 78 varieties aligned to the scaffolded Basmati 334 genome, and called the polymorphic sites segregating within this variety group. After filtering, a total of 4,430,322 SNPs across the *circum*-basmati dataset remained, which were used to examine population genetic relationships within *circum*-basmati.

We conducted principal component analysis (PCA) using the polymorphism data and color-coded each *circum*-basmati rice variety according to its country of origin (Figure 7A). The PCA suggested that *circum*-basmati rices could be divided into three major groups with clear geographic associations: (Group 1) a largely Bhutan/Nepal-based group, (Group 2) an India/Bangladesh/Myanmar-based group, and (Group 3) an Iran/Pakistan-based group. The rice varieties that could not be grouped occupied an ambiguous space across the principal components, suggesting these might represent admixed rice varieties.

**Figure 7.**
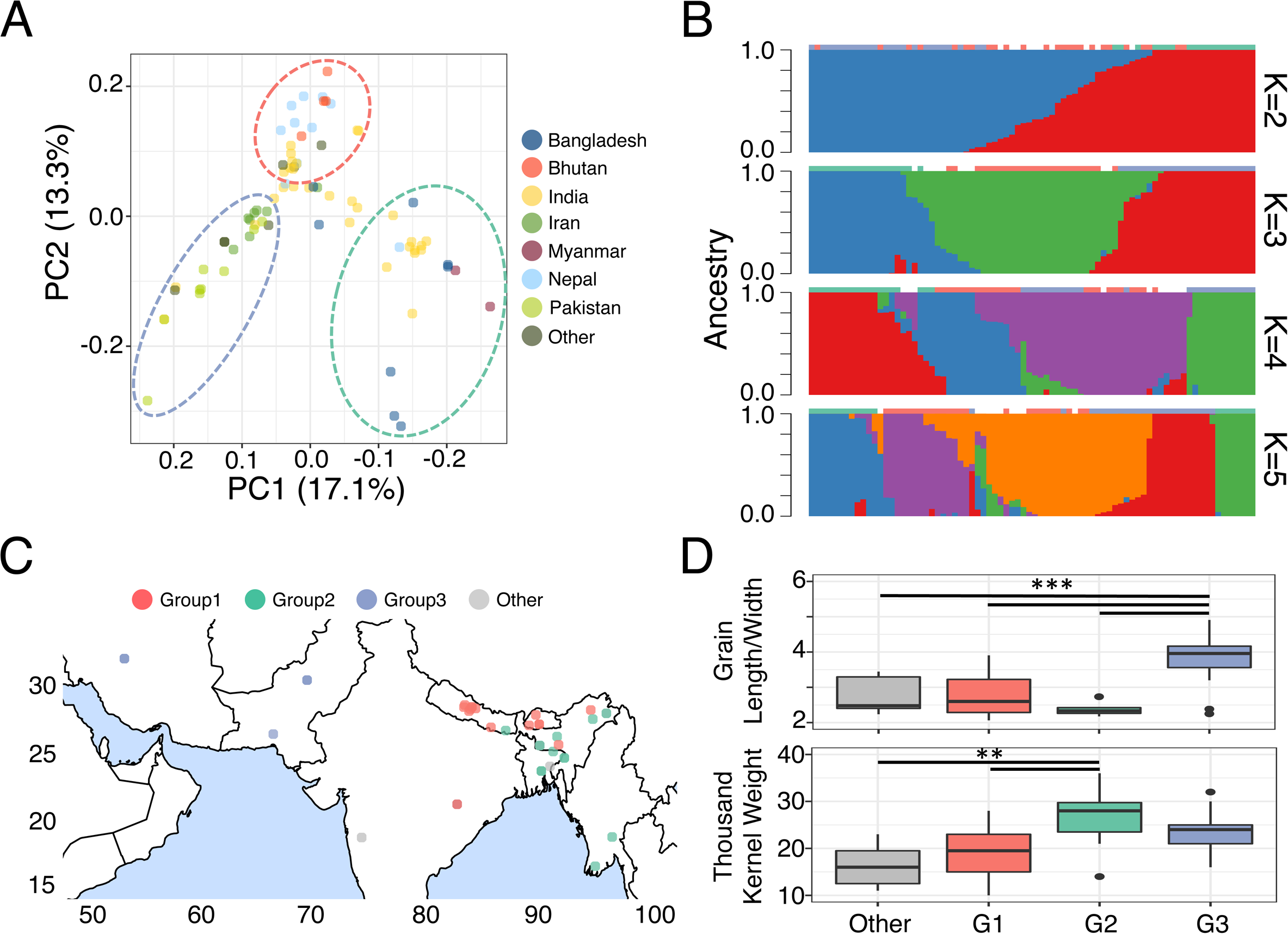
Population structure within the *circum*-basmati rices. (A) PCA plot for the 78-variety *circum*-basmati rice population genomic dataset. The three genetic groups designated by this study can be seen in the color-coded circles with dashed lines. (B) *ADMIXTURE* plot of K=2, 3, 4, and 5 for the 78 landraces. The color-coding from (A) is indicated above each sample’s ancestry proportion. (C) Geographic distribution of the 78 *circum*-basmati rice varieties with their grouping status color-coded according to (A). (D) Agronomic measurements for the 78 *circum*-basmati rice varieties sorted into the three groups designated by this study. ** indicate p-value < 0.01 and *** indicate p-value < 0.001.

To obtain better insight into the ancestry of each rice variety, we used *fastSTRUCTURE* [91] and varied assumed ancestral population (K) from 2 to 5 groups so the ancestry proportion of each rice variety could be estimated (Figure 7B). At K=2, the India/Bangladesh/Myanmar and Iran/Pakistan rice groups were shown to have distinct ancestral components, while the Bhutan/Nepal group was largely an admixture of the other two groups. At K=3, the grouping status designated from the PCA was largely concordant with the ancestral components. At K=4, most India/Bangladesh/Myanmar rices had a single ancestral component, but Iran/Pakistan rices had two ancestral components that were shared with several Bhutan/Nepal landraces. Furthermore, several of the cultivars from the latter group seemed to form an admixed group with India/Bangladesh/Myanmar varieties. In fact, when a phylogenetic tree was reconstructed using the polymorphic sites, varieties within the India/Bangladesh/Myanmar and Iran/Pakistan groups formed a monophyletic clade with each other. On the other hand, Bhutan/Nepal varieties formed a paraphyletic group where several clustered with the Iran/Pakistan varieties (Supplemental Figure 9).

In summary, the *circum*-basmati rices have evolved across a geographic gradient with at least three genetic groups (Figure 7C). These existed as distinct ancestral groups that later admixed to form several other *circum*-basmati varieties. Group 1 and Group 3 rices in particular may have experienced greater admixture, while the Group 2 landraces remained genetically more isolated from other *circum*-basmati subpopulations. We also found differences in agronomic traits associated with our designated groups (Figure 7D). The grain length to width ratio, which is a highly prized trait in certain *circum*-basmati rices [24], was significantly larger in Group 3 Iran/Pakistan varieties. The thousand-kernel weights, on the other hand, were highest for Group 2 India/Bangladesh/Myanmar varieties and were significantly higher than those for the ungrouped and Group 1 Bhutan/Nepal varieties.

## DISCUSSION

Nanopore sequencing is becoming an increasingly popular approach to sequence and assemble the often large and complex genomes of plants [92–94]. Here, using long-read sequences generated with Oxford Nanopore Technologies’ sequencing platform, we assembled genomes of two *circum*-basmati rice cultivars, with quality metrics that were comparable to other rice variety group reference genome assemblies [37, 40, 41]. With modest genome coverage, we were able to develop reference genome assemblies that represented a significant improvement over a previous *circum*-basmati reference genome sequence, which had been assembled with a > 3-fold higher genome coverage than ours, but from short-read sequences [42]. With additional short-read sequencing reads, we were able to correct errors from the nanopore sequencing reads, resulting in two high-quality *circum*-basmati genome assemblies.

Even with long-read sequence data, developing good plant reference genome sequences still requires additional technologies such as optical mapping or Hi-C sequencing for improving assembly contiguity [95–98], which can be error prone as well [56]. Our assemblies were also fragmented into multiple contigs, but sizes of these contigs were sufficiently large that we could use reference genome sequences from another rice variety group to anchor the majority of contigs and scaffold them to higher-order chromosome-level assemblies. Hence, with a highly contiguous draft genome assembly, reference genome-based scaffolding can be a cost-efficient and powerful method of generating chromosome-level assemblies.

Repetitive DNA constitutes large proportions of plant genomes [99], and there is an advantage to using long-read sequences for genome assembly as it enables better annotation of transposable elements. Many transposable element insertions have evolutionarily deleterious consequences in the rice genome [54, 100, 101], but some insertions could have beneficial effects on the host [102]. Using our genome assembly, we have identified retrotransposon families that have expanded specifically within *circum*-basmati genomes. While more study will be necessary to understand the functional effects of these insertions, long-read sequences have greatly improved the assembly and identification of repeat types.

Due to a lack of archaeobotanical data, the origins of *circum*-basmati rice have remained elusive. Studies of this variety group’s origins have primarily focused on genetic differences that exist between *circum*-basmati and other Asian rice variety groups [6, 7]. Recently, a study suggested that *circum*-basmati rice (called ‘aromatic’ in that study) was a product of hybridization between the *circum*-aus and japonica rice variety groups [17]. This inference was based on observations of phylogenetic relationships across genomic regions that showed evidence of domestication-related selective sweeps. These regions mostly grouped *circum*-basmati with japonica or *circum*-aus. In addition, chloroplast haplotype analysis indicated that most *circum*-basmati varieties carried a chloroplast derived from a wild rice most closely related to *circum*-aus landraces [103]. Our evolutionary analysis of *circum*-basmati rice genomes generally supported this view. Although our results suggest that *circum*-basmati had its origins primarily in japonica, we also find significant evidence of gene flow originating from *circum*-aus, which we detected both in comparative genomic and population genomic analyses. Demographic modeling indicated a period of isolation among *circum*-aus, *circum*-basmati, and japonica, with gene flow occurring only after lineage splitting of each group. Here, our model is consistent with the current view that gene flow is a key evolutionary process associated with the diversification of rice [10, 12–14, 16, 104, 105].

Interestingly, we found that chromosome 10 of *circum*-basmati had an evolutionary history that differed significantly from that of other chromosomes. Specifically, compared to japonica, this chromosome had the highest proportion of presence/absence variation, and shared more alleles with *circum*-aus. Based on this result, we hypothesize that this is largely due to higher levels of introgression from *circum*-aus into chromosome 10 compared to other chromosomes. Such a deviation of evolutionary patterns on a single chromosome has been observed in the *Aquilegia* genus [106], but to our knowledge has not been observed elsewhere. Why this occurred is unclear at present, but it may be that selection has driven a higher proportion of *circum*-aus alleles into chromosome 10. Future work will be necessary to clarify the consequence of this higher level of admixture on chromosome 10.

Very little is known about population genomic diversity within *circum*-basmati. Our analysis suggests the existence of at least three genetic groups within this variety group, and these groups showed geographic structuring. Several varieties from Group 1 (Bhutan/Nepal) and Group 3 (Iran/Pakistan) had population genomic signatures consistent with an admixed population, while Group 2 (India/Bangladesh/Myanmar) was genetically more distinct from the other two subpopulations. In addition, the geographic location of the India/Bangladesh/Myanmar group largely overlaps the region where *circum*-aus varieties were historically grown [107, 108]. Given the extensive history of admixture that *circum*-basmati rices have with *circum*-aus, the India/Bangladesh/Myanmar group may have been influenced particularly strongly by gene flow from *circum*-aus. How these three genetic subpopulations were established may require a deeper sampling with in-depth analysis, but the geographically structured genomic variation shows that the diversity of *circum*-basmati has clearly been underappreciated. In addition, the Basmati 334 and Dom Sufid varieties, for which we generated genome assemblies in this study, both belong to the Iran/Pakistan genetic group. Thus, our study still leaves a gap in our knowledge of genomic variation in the Bhutan/Nepal and India/Bangladesh/Myanmar genetic groups, and varieties in these groups would be obvious next targets for generating additional genome assemblies.

## CONCLUSIONS

In conclusion, our study shows that generating high-quality plant genome assemblies is feasible with relatively modest amounts of resources and data. Using nanopore sequencing, we were able to produce contiguous, chromosome-level genome assemblies for cultivars in a rice variety group that contains economically and culturally important varieties. Our reference genome sequences have the potential to be important genomic resources for identifying single nucleotide polymorphisms and larger structural variations that are unique to *circum*-basmati rice. Analyzing *de novo* genome assemblies for a larger sample of Asian rice will be important for uncovering and studying hidden population genomic variation too complex to study with only short-read sequencing technology.

## MATERIALS AND METHODS

### Plant material

Basmati 334 (IRGC 27819; GeneSys passport: https://purl.org/germplasm/id/23601903-f8c3-4642-a7fc-516a5bc154f7) is a basmati (*sensu stricto*) landrace from Pakistan and was originally donated to the International Rice Research Institute (IRRI) by the Agricultural Research Council (ARC) in Karachi (donor accession ID: PAK. SR. NO. 39). Dom Sufid (IRGC 117265; GeneSys passport: https://purl.org/germplasm/id/fb861458-09de-46c4-b9ca-f5c439822919) is a sadri landrace from Iran. Seeds from accessions IRGC 27819 and IRGC 117265 were obtained from the IRRI seed bank, surface-sterilized with bleach, and germinated in the dark on a wet paper towel for four days. Seedlings were transplanted individually in pots containing continuously wet soil in a greenhouse at New York University’s Center for Genomics and Systems Biology and cultivated under a 12h day-12h night photoperiod at 30°C. Plants were kept in the dark in a growth cabinet under the same climatic conditions for four days prior to tissue harvesting. Continuous darkness induced chloroplast degradation, which diminishes the amount of chloroplast DNA that would otherwise end up in the DNA extracted from the leaves.

### DNA extractions

Thirty-six 100-mg samples (3.6 g total) of leaf tissue from a total of 10 one-month-old plants were flash-frozen at harvest for each accession and stored at −80°C. DNA extractions were performed by isolating the cell nuclei and gently lysing the nuclei to extract intact DNA molecules [109]. Yields ranged between 140ng/ul and 150ng/ul.

### Library preparation and nanopore sequencing

Genomic DNA was visualized on an agarose gel to determine shearing. DNA was size-selected using BluePippin BLF7510 cassette (Sage Science) and high-pass mode (>20 kb) and prepared using Oxford Nanopore Technologies’ standard ligation sequencing kit SQK-LSK109. FLO-MIN106 (R9.4) flowcells were used for sequencing on the GridION X5 platform.

### Library preparation and Illumina sequencing

Extracted genomic DNA was prepared for short-read sequencing using the Illumina Nextera DNA Library Preparation Kit. Sequencing was done on the Illumina HiSeq 2500 – HighOutput Mode v3 with 2×100 bp read configuration, at the New York University Genomics Core Facility.

### Genome assembly, polishing, and scaffolding

After completion of sequencing, the raw signal intensity data was used for base calling using *flip flop* (version 2.3.5) from Oxford Nanopore Technologies. Reads with a mean qscore (quality) greater than 8 and a read length greater than 8 kb were used, and trimmed for adaptor sequences using *Porechop* (https://github.com/rrwick/Porechop). Raw nanopore sequencing reads were corrected using the program *Canu* [110], and then assembled with the genome assembler *Flye* [111].

The initial draft assemblies were polished for three rounds using the raw nanopore reads with *Racon* ver. 1.2.1 [112], and one round with *Medaka* (https://github.com/nanoporetech/medaka) from Oxford Nanopore Technologies. Afterwards, reads from Illunima sequencing were used by *bwa-mem* [113] to align to the draft genome assemblies. The alignment files were then used by *Pilon* ver. 1.22 [114] for three rounds of polishing.

Contigs were scaffolded using a reference genome-guided scaffolding approach implemented in *RaGOO* [56]. Using the Nipponbare genome as a reference, we aligned the *circum*-basmati genomes using *Minimap2* [115]. *RaGOO* was then used to order the assembly contigs. Space between contigs was artificially filled in with 100 ‘N’ blocks.

Genome assembly statistics were calculated using the *bbmap stats.sh* script from the *BBTools* suite (https://jgi.doe.gov/data-and-tools/bbtools/). Completeness of the genome assemblies was evaluated using *BUSCO* ver. 2.0 [116]. Synteny between the *circum*-basmati genomes and the Nipponbare genome was visualized using *D-GENIES* [117]. Genome-wide dotplot from *D-GENIES* indicated the initial genome assembly of Dom Sufid had an evidence of a large chromosomal fusion between the ends of chromosome 4 and 10. Closer examination of this contig (named contig_28 of Dom Sufid) showed the break point overlapped the telomeric repeat sequence, indicating there had been a misassembly between the ends of chromosome 4 and 10. Hence, contig_28 was broken up into two so that each contig represented the respective chromosome of origin, and were then subsequently scaffolded using *RaGOO*.

Inversions that were observed in the dot plot were computationally verified independently using raw nanopore reads. The long read-aware aligner *ngmlr* [55] was used to align the nanopore reads to the Nipponbare genome, after which the long read-aware structural variation caller *sniffles* [55] was used to call and detect inversions.

The number of sites aligning to the Nipponbare genome was determined using the *Mummer4* package [118]. Alignment delta files were analyzed with the *dnadiff* suite from the *Mummer4* package to calculate the number of aligned sites, and the number of differences between the Nipponbare genome and the *circum*-basmati genomes.

### Gene annotation and analysis

Gene annotation was conducted using the *MAKER* program [52, 53]. An in-depth description of running *MAKER* can be found on the website: https://gist.github.com/darencard/bb1001ac1532dd4225b030cf0cd61ce2. We used published *Oryza* genic sequences as evidence for the gene modeling process. We downloaded the Nipponbare cDNA sequences from RAP-DB (https://rapdb.dna.affrc.go.jp/) to supply as EST evidence, while the protein sequences from the 13 *Oryza* species project [37] were used as protein evidence for the *MAKER* pipeline. Repetitive regions identified from the repeat analysis were used to mask out the repeat regions for this analysis. After a first round of running *MAKER* the predicted genes were used by *SNAP* [119] and *Augustus* [120] to create a training dataset of gene models, which was then used for a second round of *MAKER* gene annotation.

Orthology between the genes from different rice genomes was determined with *Orthofinder* ver. 1.1.9 [59]. Ortholog statuses were visualized with the *UpSetR* package [121].

Gene ontology for the orthogroups that are missing specifically in the *circum*-basmati were examined by using the japonica Nipponbare gene, and conducting a gene ontology enrichment analysis on *agriGO* v2.0 [122]. Gene ontology enrichment analysis for the *circum*-basmati specific orthogroups was conducted first by predicting the function and gene ontology of each *circum*-basmati genome gene model using the eggnog pipeline [123]. We required an ontology to have more than 10 genes as a member for further consideration, and enrichment was tested through a hypergeometric test using the *GOstat* package [124].

### Repetitive DNA annotation

The repeat content of each genome assembly was determined using *Repeatmasker* ver. 4.0.5 (http://www.repeatmasker.org/RMDownload.html). We used the *Oryza*-specific repeat sequences that were identified from Choi et al. [14] (DOI: 10.5061/dryad.7cr0q), who had used *Repeatmodeler* ver. 1.0.8 (http://www.repeatmasker.org/RepeatModeler.html) to *de novo*-annotate repetitive elements across wild and domesticated *Oryza* genomes [37].

LTR retrotransposons were annotated using the program *LTRharvest* [125] with parameters adapted from [126]. LTR retrotransposons were classified into superfamilies [76] using the program *RepeatClassifier* from the *RepeatModeler* suite. Annotated LTR retrotransposons were further classified into specific families using the 242 consensus sequences of LTR-RTs from the RetrOryza database [83]. We used *blastn* [127] to search the RetrOryza sequences, and each of our candidate LTR retrotransposons was identified using the “80-80-80” rule [76]: two TEs belong to the same family if they were 80% identical over at least 80 bp and L 80% of their length.

Insertion times for the LTR retrotransposons were estimated using the DNA divergence between pairs of LTR sequences [75]. The L-INS-I algorithm in the alignment program *MAFFT* ver. 7.154b [128] was used to align the LTR sequences. *PAML* ver. 4.8 [129] was used to estimate the DNA divergence between the LTR sequences with the Kimura-2-parameter base substitution model [130]. DNA divergence was converted to divergence time (i.e. time since the insertion of a LTR retrotransposon) approximating a base substitution rate of 1.3×10^-8^ [131], which is two times higher than the synonymous site substitution rate.

### Presence/absence variation detection

PAVs between the Nipponbare genome and the *circum*-Basmati assemblies were detected using the *Assemblytics* suites [60]. Initially, the Nipponbare genome was used as the reference to align the *circum*-basmati assemblies using the program *Minimap2*. The resulting SAM files were converted to files in delta format using the *sam2delta.py* script from the *RaGOO* suite. The delta files were then uploaded onto the online *Assemblytics* analysis pipeline (http://assemblytics.com/). Repetitive regions would cause multiple regions in the Nipponbare or *circum*-basmati genomes to align to one another, and in that case *Assemblytics* would call the same region as a PAV multiple times. Hence, any PAV regions that overlapped for at least 70% of their genomic coordinates were collapsed to a single region.

The combination of *ngmlr* and *sniffles* was also used to detect the PAVs that differed between the Nipponbare genome and the raw nanopore reads for the *circum*-basmati rices. Because *Assemblytics* only detects PAVs in the range of 50 bp to 100,000 bp, we used this window as a size limit to filter out the PAVs called by *sniffles*. Only PAVs supported by more than 5 reads by *sniffles* were analyzed.

*Assemblytics* and sniffles call the breakpoints of PAVs differently. *Assemblytics* calls a single-best breakpoint based on the genome alignment, while *sniffles* calls a breakpoint across a predicted interval. To find overlapping PAVs between *Assemblytics* and *sniffles* we added 500 bp upstream and downstream of the *Assemblytics*-predicted breakpoint positions.

### Detecting gene deletions across the *circum-*basmati population

Genome-wide deletion frequencies of each gene were estimated using the 78-variety *circum*-basmati population genomic dataset. For each of the 78 varieties, raw sequencing reads were aligned to the *circum*-basmati and Nipponbare genomes using *bwa-mem*. Genome coverage per site was calculated using *bedtools genomecov* [132]. For each variety the average read coverage was calculated for each gene, and a gene was designated as deleted if its average coverage was less than 0.05×.

### Whole-genome alignment of *Oryza* genomes assembled *de novo*

Several genomes from published studies that were assembled *de novo* were analyzed. These include domesticated Asian rice genomes from the japonica variety group cv. Nipponbare [33]; the indica variety group cvs. 93-11 [32], IR8 [37], IR64 [38], MH63 [40], R498 [41], and ZS97 [40]; the *circum*-aus variety group cvs. DJ123 [38], Kasalath [39], and N22 [37]; and the *circum*-basmati variety group cv. GP295-1 [42]. Three genomes from wild rice species were also analyzed; these were *O. barthii* [35], *O. nivara* [37], and *O. rufipogon* [37].

Alignment of the genomes assembled *de novo* was conducted using the approach outlined in Haudry *et al.* [133], and the alignment has been used in another rice comparative genomic study [14]. Briefly, this involved using the Nipponbare genome as the reference for aligning all other genome assemblies. Alignment between japonica and a query genome was conducted using *LASTZ* ver. 1.03.73 [134], and the alignment blocks were chained together using the UCSC Kent utilities [135]. For japonica genomic regions with multiple chains, the chain with the highest alignment score was chosen as the single-most orthologous region. This analyzes only one of the multiple regions that are potentially paralogous between the japonica and query genomes, but this was not expected to affect the downstream phylogenomic analysis of determining the origin and evolution of the *circum-*basmati rice variety group. All pairwise genome alignments between the japonica and query genomes were combined into a multi-genome alignment using *MULTIZ* [136].

### Phylogenomic analysis

The multi-genome alignment was used to reconstruct the phylogenetic relationships between the domesticated and wild rices. Four-fold degenerate sites based on the gene model of the reference japonica genome were extracted using the *msa_view* program from the *phast* package ver. 1.4 [137]. The four-fold degenerate sites were used by *RAxML* ver. 8.2.5 [138] to build a maximum likelihood-based tree, using a general time-reversible DNA substitution model with gamma-distributed rate variation.

To investigate the genome-wide landscape of introgression and incomplete lineage sorting we examined the phylogenetic topologies of each gene [139]. For a three-species phylogeny using *O. barthii* as an outgroup there are three possible topologies. For each gene, topology-testing methods [140] can be used to determine which topology significantly fits the gene of interest [14]. *RAxML*-estimated site-likelihood values were calculated for each gene and the significant topology was determined using the Approximately Unbiased (AU) test [86] from the program *CONSEL* v. 0.20 [141]. Genes with AU test results with a likelihood difference of zero were omitted and the topology with an AU test support of greater than 0.95 was selected.

### Testing for evidence of admixture

Evidence of admixture between variety groups was detected using the ABBA-BABA test D-statistics [87, 88]. In a rooted three-taxon phylogeny [i.e. “((P1,P2),P3),O” where P1, P2, and P3 are the variety groups of interest and O is outgroup *O. barthii*], admixture can be inferred from the combination of ancestral (“A”) and derived (“B”) allelic states of each individual. The ABBA conformation arises when variety groups P2 and P3 share derived alleles, while the BABA conformation is found when P1 and P3 share derived alleles. The difference in the frequency of the ABBA and BABA conformations is measured by the D-statistics, where significantly positive D-statistics indicate admixture between the P2 and P3 variety groups, and significantly negative D-statistics indicate admixture between the P1 and P3 variety groups. The genome was divided into 100,000-bp bins for jackknife resampling and calculation of the standard errors. The significance of the D-statistics was calculated using the Z-test, and D-statistics with Z-scores greater than |3.9| (p < 0.0001) were considered significant.

### Population genomic analysis

We downloaded FASTQ files from the 3K Rice Genome Project [7] for rice varieties that were determined to be *circum*-basmati varieties in that project. An additional 8 *circum*-basmati varieties were sequenced on the Illumina sequencing platform as part of this study. The raw reads were aligned to the scaffolded Basmati 334 genome using the program *bwa-mem*. PCR duplicates were determined computationally and removed using the program *picard* version 2.9.0 (http://broadinstitute.github.io/picard/). Genotype calls for each site were conducted using the *GATK HaplotypeCaller* engine using the option ‘-ERC GVCF’. The output files were in the genomic variant call format (gVCF), and the gVCFs from each variety were merged using the *GATK GenotypeGVCFs* engine.

SNP and INDEL variants from the population variant file were filtered independently using the *GATK* bestpractice hard filter pipeline [142]. SNP variants within 5 bps of an INDEL variant were filtered. *Vcftools* version 0.1.15 [143] was used to filter sites for which genotypes were not called for more than 20% of the varieties Because domesticated rice is an inbreeding species we also implemented a heterozygosity filter by filtering out sites that had a heterozygote genotype in more than 5% of the samples using the program *vcffilterjdk.jar* from the *jvarkit* suite (https://figshare.com/articles/JVarkit_java_based_utilities_for_Bioinformatics/1425030). Missing genotypes were imputed and phased using *Beagle* version 4.1 [144].

To examine the within-*circum*-basmati variety group population structure we first randomly pruned the sites by sampling a polymorphic site every 200,000 bp using *plink* [145]. *Plink* was also used to conduct a principal component analysis. Ancestry proportions of each sample were estimated using *fastSTRUCTURE* [91]. A neighbor-joining tree was built by calculating the pairwise genetic distances between samples using the Kronecker delta function-based equation [146]. From the genetic distance matrix a neighbor-joining tree was built using the program *FastME* [147].

### Evolutionary relationships among the *circum-*basmati, *circum-*aus, and japonica populations

To investigate the evolutionary origins of the *circum*-basmati population, we focused on the landrace varieties that had been sequenced with a genome-wide coverage of greater than 14×. The population data for the *circum*-aus and japonica populations were obtained from the 3K Rice Genome Project [7], from which we also analyzed only the landrace varieties that had been sequenced with a genome-wide coverage greater than 14×. For an outgroup, we obtained *O. barthii* sequencing data from previous studies [35, 148], and focused on the samples that were not likely to be feralized rices [148]. The Illumina reads were aligned to the scaffolded Basmati 334 genome and SNPs were called and filtered according to the procedure outlined in the “Population genomic analysis” section.

We examined the genome-wide local topological relationship using *twisst* [89]. Initially, a sliding window analysis was conducted to estimate the local phylogenetic trees in windows with a size of 100 or 500 polymorphic sites using *RAxML* with the GTRCAT substitution model. The script *raxml_sliding_windows.py* from the *genomics_general* package by Simon Martin (https://github.com/simonhmartin/genomics_general/tree/master/phylo) was used. The ‘complete’ option of *twisst* was used to calculate the exact weighting of each local window.

### □a□i demographic model

The demography model underlying the evolution of *circum*-basmati D D rice was tested using the diffusion approximation method of *δaδi* [90]. A visual representation of the 13 demographic models that were examined can be seen in Supplementary Figure S6. The population group and genotype calls used in the twisst analysis were also used to calculate the site allele frequencies. Polymorphic sites were polarized using the *O. barthii* reference genome. We used a previously published approach [148], which generates an *O. barthii*-ized basmati genome sequence. This was accomplished using the Basmati 334 reference genome to align the *O. barthii* genome. For every basmati genome sequence position was then changed into the aligned *O. barthii* sequence. Gaps, missing sequence, and repetitive DNA region were denoted as ‘N’.

We optimized the model parameter estimates using the Nelder-Mead method and randomly perturbed the parameter values for four rounds. Parameter values were perturbed for three-fold, two-fold, two-fold, and one-fold in each subsequent round, while the perturbation was conducted for 10, 20, 30, and 40 replicates in each subsequent round. In each round parameter values from the best likelihood model of the previous round were used as the starting parameter values for the next round. Parameter values from the round with the highest likelihood were chosen to parameterize each demographic model. Akaike Information Criteria (AIC) values were used to compare demography models. The demography model with the lowest AIC was chosen as the best-fitting model.

### Agronomic trait measurements

Data on geolocation of collection as well as on seed dimensions and seed weight for each of the *circum*-basmati landrace varieties included in this study were obtained from passport data included in the online platform Genesys (https://www.genesys-pgr.org/welcome).

## DECLARATIONS

### Ethics approval and consent to participate

Not applicable.

### Consent for publication

Not applicable.

### Availability of data and materials

Raw nanopore sequencing FAST5 files generated from this study are available at the European Nucleotide Archive under bioproject ID PRJEB28274 (ERX3327648-ERX3327652) for Basmati 334 and PRJEB32431 (ERX3334790-ERX3334793) for Dom Sufid. Associated FASTQ files are available under ERX3498039-ERX3498043 for Basmati 334 and ERX3498024-ERX3498027 for Dom Sufid. Illumina sequencing generated from this study can be found under bioproject ID PRJNA422249 and PRJNA557122. A genome browser for both genome assemblies can be found at http://purugganan-genomebrowser.bio.nyu.edu/cgi-bin/hgTracks?db=Basmati334 for Basmati 334, and http://purugganan-genomebrowser.bio.nyu.edu/cgi-bin/hgTracks?db=DomSufid for Dom Sufid. All data including the assembly, annotation, genome alignment, and population VCFs generated from this study can be found at https://doi.org/10.5281/zenodo.3355330.

### Competing interests

XD, PR, EDH, and SJ are employees of Oxford Nanopore Technologies and are shareholders and/or share option holders.

### Funding

This work was supported by grants from the Gordon and Betty Moore Foundation through Grant GBMF2550.06 to S.C.G., and from the National Science Foundation Plant Genome Research Program (IOS-1546218), the Zegar Family Foundation (A16-0051) and the NYU Abu Dhabi Research Institute (G1205) to M.D.P. The funding body had no role in the design of the study and collection, analysis, and interpretation of data and in writing the manuscript.

### Authors’ contributions

JYC, SCG, SZ, and MDP conceived the project and its components. JYC, SCG, and SZ prepared the sample material for sequencing. XD, PR, EDH, and SJ conducted the genome sequencing and assembling. JYC, ZNL, and SCG performed the data analysis. JYC and ZNL prepared the figures and tables. JYC and MDP wrote the manuscript with help from ZNL and SCG.

## Acknowledgements

We thank Katherine Dorph for assistance with growing and maintaining the plants, and Adrian Platts for computational support. We thank Rod Wing, David Kudrna, and Jayson Talag from Arizona Genomics Institute with the high-molecular weight DNA extraction. We thank the New York University Genomics Core Facility for sequencing support and the New York University High Performance Computing for supplying the computational resources.

## Supplemental Figures

Supplemental Figure 1. Dot plot comparing chromosome 6 of japonica variety Nipponbare to *circum*-aus variety N22 and indica variety R498.

Supplemental Figure 2. Distribution of the proportion of missing nucleotides for japonica variety Nipponbare gene models across the orthologous non-japonica genomic regions.

Supplemental Figure 3. Effect of coverage threshold to call a deletion and the total number of deletion calls for samples with various genome coverage.

**Supplemental Figure 4. Insertion time of LTR retrotransposon in various *Oryza* variety group genomes**. Number of annotated LTR retrotransposons is shown above boxplot. The variety group genomes that do not have a significantly different insertion time after a Tukey’s range test are indicated with the same letter.

Supplemental Figure 5. Density of presence-absence variation (PAV) per 500,000 bp window for each chromosome.

**Supplemental Figure 6. Genome-wide topology weight from 500 SNP size window**. Chromosomal distribution of topology weights involving trios of the *circum*-aus, *circum*-basmati, and japonica rices (left), and the sum of the topology weights (right).

Supplemental Figure 7. 13 demographic models tested by □a□i.

**Supplemental Figure 8. □a□i model fit for the best-fitting demographic model.** Above row shows the observed and model fit folded site frequency spectrum. Below shows the map and histogram of the residuals.

Supplemental Figure 9. Neighbor-joining phylogenetic tree of the 78 *circum*-basmati population sample.

Supplemental Table 1. Inversion detect by *sniffles* in the Nipponbare reference genome.

Supplemental Table 2. The 78 *circum*-basmati samples with Illumina sequencing result used in this study.

Supplemental Table 3. Names of the Basmati 334 and Dom Sufid genome gene models that had a deletion frequency of zero across the population.

Supplemental Table 4. Names of the Basmati 334 and Dom Sufid genome gene models that had a deletion frequency of above 0.3 and omitted from down stream analysis.

Supplemental Table 5. Orthogroup status for the Basmati 334, Dom Sufid, R498, Nipponbare, and N22 genome gene models.

Supplemental Table 6. Count and repeat types of the presence-absence variation (PAV) in the Basmati 334 or Dom Sufid genome in comparison to the Nipponbare genome.

Supplemental Table 7. Gene ontology results for orthogroups where gene members from the *circum*-basmati are missing.

Supplemental Table 8. Gene ontology results for orthogroups where gene members from

*circum*-aus, indica, and japonica are missing.

Supplemental Table 9. Population frequency across the 78 *circum*-basmati samples for orthogroups that were specifically missing a gene in the Basmati 334 and Dom Sufid genome gene models.

Supplemental Table 10. Genome coordinates of the LTR retrotransposons of the Basmati 334 genomes.

Supplemental Table 11. Genome coordinates of the LTR retrotransposons of the Dom Sufid genomes.

Supplemental Table 12. Genome coordinates of the Gypsy elements indicated with a single star in Figure 3.

Supplemental Table 13. Genome coordinates of the Copia elements indicated with a single star in Figure 3.

Supplemental Table 14. Genome coordinates of the Gypsy elements indicated with a double star in Figure 3.

Supplemental Table 15. Genome coordinates of the Copia elements indicated with a triple star in Figure 3.

Supplemental Table 16. The 82 *Oryza* population samples with Illumina sequencing result used in this study.

**Supplemental Table 17. DaDi parameter estimates for the 13 different demographic models**. See supplemental figure 7 for visualization of the estimating parameters.

## Notes

#### Summary of Updates

New results and major analysis done in new version

DOI:10.5281/zenodo.3355330

