## Supplementary material for "Nanopore-based genome assembly and the evolutionary genomics of basmati rice": Supplemental Figure 4.pptx

### Slide 1
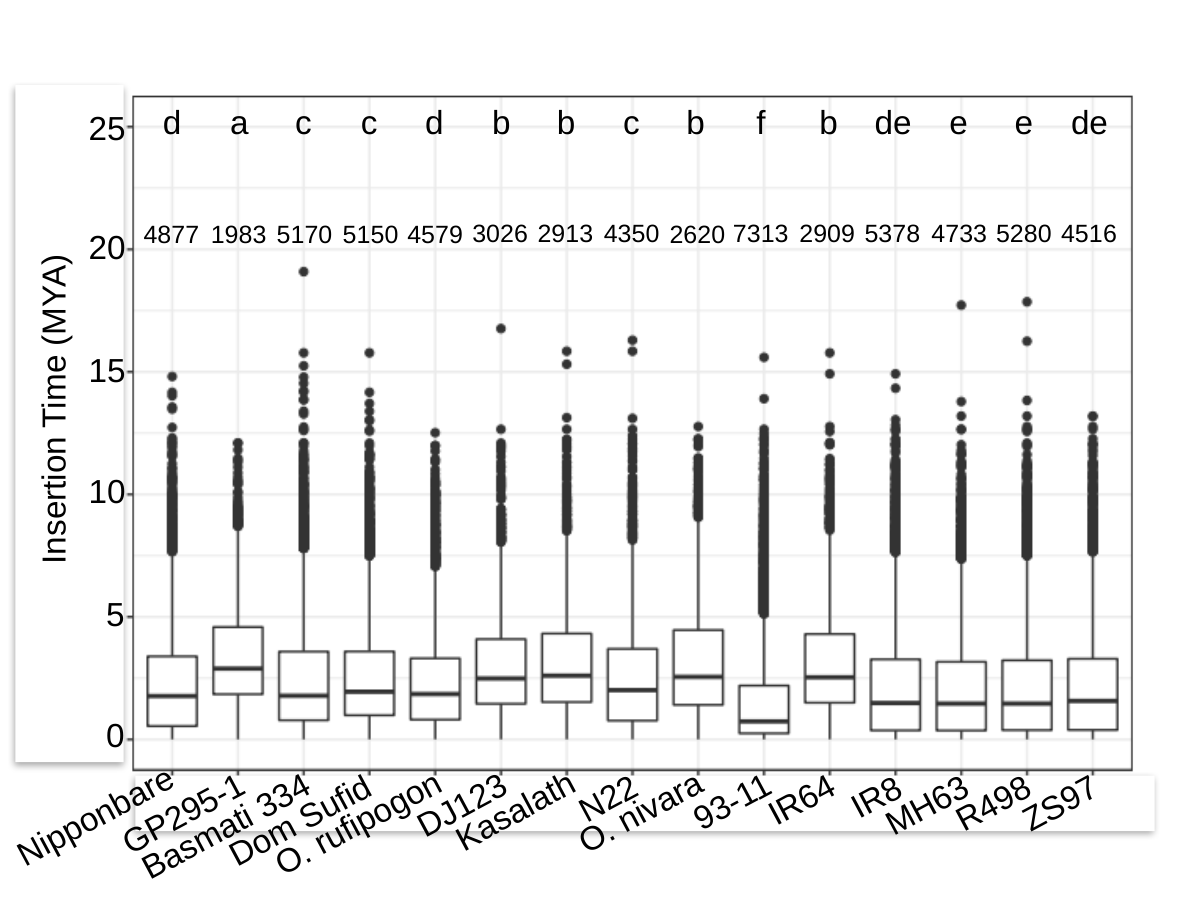

d
a
c
c
d
b
b
c
b
f
b
de
e
e
de
25
3026
2913
4350
7313
2909
5378
4733
5280
4516
4877
1983
2620
5170
5150
4579
20
15
Insertion Time (MYA)
10
5
0
IR8
N22
IR64
93-11
R498
ZS97
MH63
DJ123
Kasalath
O. nivara
GP295-1
Nipponbare
Dom Sufid
O. rufipogon
Basmati 334
