## Supplementary material for "Nanopore-based genome assembly and the evolutionary genomics of basmati rice": Supplemental Figure 6.pptx

### Slide 1
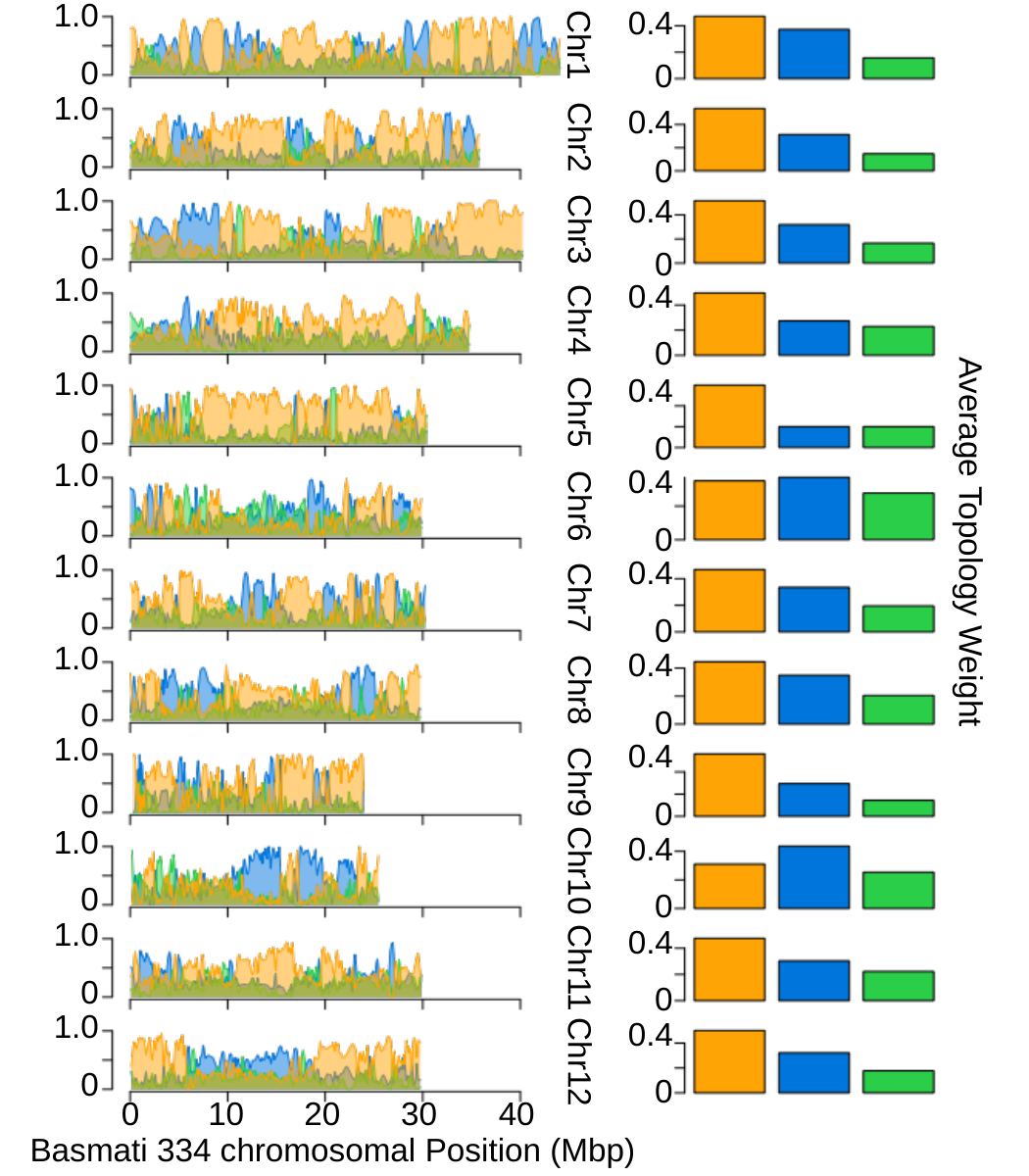

1.0
 0.4
Chr1
 0
 0
 1.0
 0.4
Chr2
 0
 0
 1.0
 0.4
Chr3
 0
 0
 1.0
 0.4
Chr4
 0
 0
 1.0
 0.4
Chr5
 0
 0
 1.0
 0.4
Chr6
 0
 0
Average Topology Weight
 1.0
 0.4
Chr7
 0
 0
 1.0
 0.4
Chr8
 0
 0
 1.0
 0.4
Chr9
 0
 0
 1.0
 0.4
Chr10
 0
 0
 1.0
 0.4
Chr11
 0
 0
 1.0
 0.4
Chr12
 0
 0
 0
10
20
30
40
Basmati 334 chromosomal Position (Mbp)
