## Supplementary material for "Nanopore-based genome assembly and the evolutionary genomics of basmati rice": Supplemental Figure 7.pptx

### Slide 1
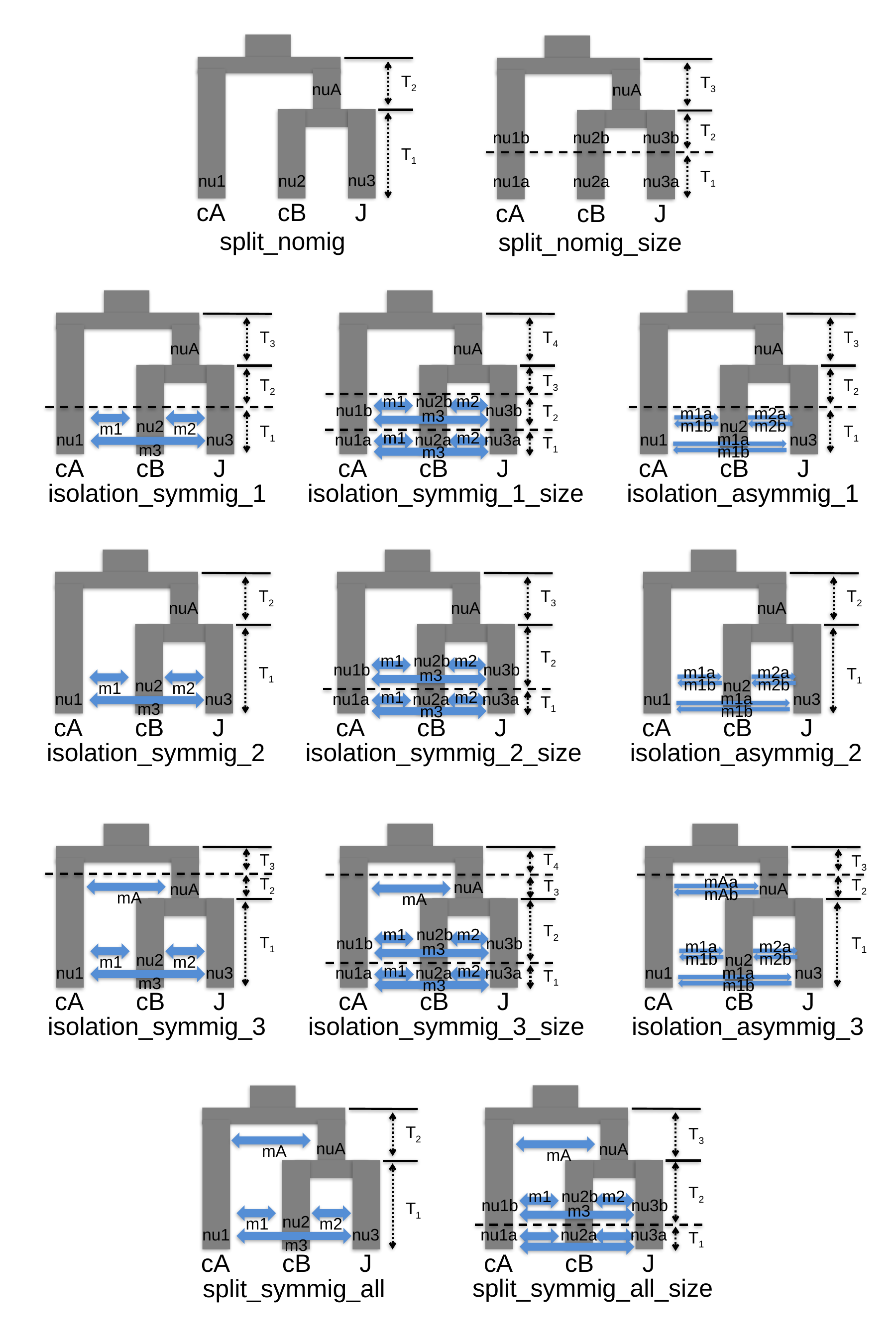

T_2_
T_1_
cA
cB
J
nuA
nu3
nu1
nu2
split_nomig
T_3_
nuA
T_2_
nu3b
nu1b
nu2b
T_1_
nu3a
nu1a
nu2a
cA
cB
J
split_nomig_size
T_3_
nuA
T_2_
nu2
m1
m2
T_1_
nu3
nu1
m3
cA
cB
J
isolation_symmig_1
T_4_
nuA
T_3_
m1
m2
nu2b
nu3b
nu1b
T_2_
m3
nu2a
nu3a
nu1a
T_1_
cA
cB
J
isolation_symmig_1_size
m1
m2
m3
T_3_
nuA
T_2_
nu2
T_1_
nu3
nu1
cA
cB
J
m1a
m2a
m1b
m2b
m1a
m1b
isolation_asymmig_1
T_2_
nuA
T_1_
nu2
m1
m2
nu3
nu1
m3
cA
cB
J
isolation_symmig_2
T_3_
nuA
T_2_
m1
m2
nu2b
nu3b
nu1b
m3
nu2a
nu3a
nu1a
T_1_
cA
cB
J
isolation_symmig_2_size
m1
m2
m3
T_2_
nuA
T_1_
nu2
nu3
nu1
cA
cB
J
m1a
m2a
m1b
m2b
m1a
m1b
isolation_asymmig_2
T_3_
T_2_
nuA
T_1_
nu2
m1
m2
nu3
nu1
m3
cA
cB
J
mA
isolation_symmig_3
T_4_
nuA
T_2_
m1
m2
nu2b
nu3b
nu1b
m3
nu2a
nu3a
nu1a
T_1_
cA
cB
J
isolation_symmig_3_size
nuA
nu2
nu3
nu1
cA
cB
J
m1a
m2a
m1b
m2b
m1a
m1b
isolation_asymmig_3
T_3_
mAa
T_2_
T_3_
mAb
mA
T_1_
m1
m2
m3
T_3_
nuA
T_2_
m1
m2
nu2b
nu3b
nu1b
m3
nu2a
nu3a
nu1a
T_1_
cA
cB
J
split_symmig_all_size
T_2_
nuA
T_1_
nu2
m1
m2
nu3
nu1
m3
cA
cB
J
split_symmig_all
mA
mA
