## Supplementary material for "Nanopore-based genome assembly and the evolutionary genomics of basmati rice": Supplemental Figure 8.pptx

### Slide 1
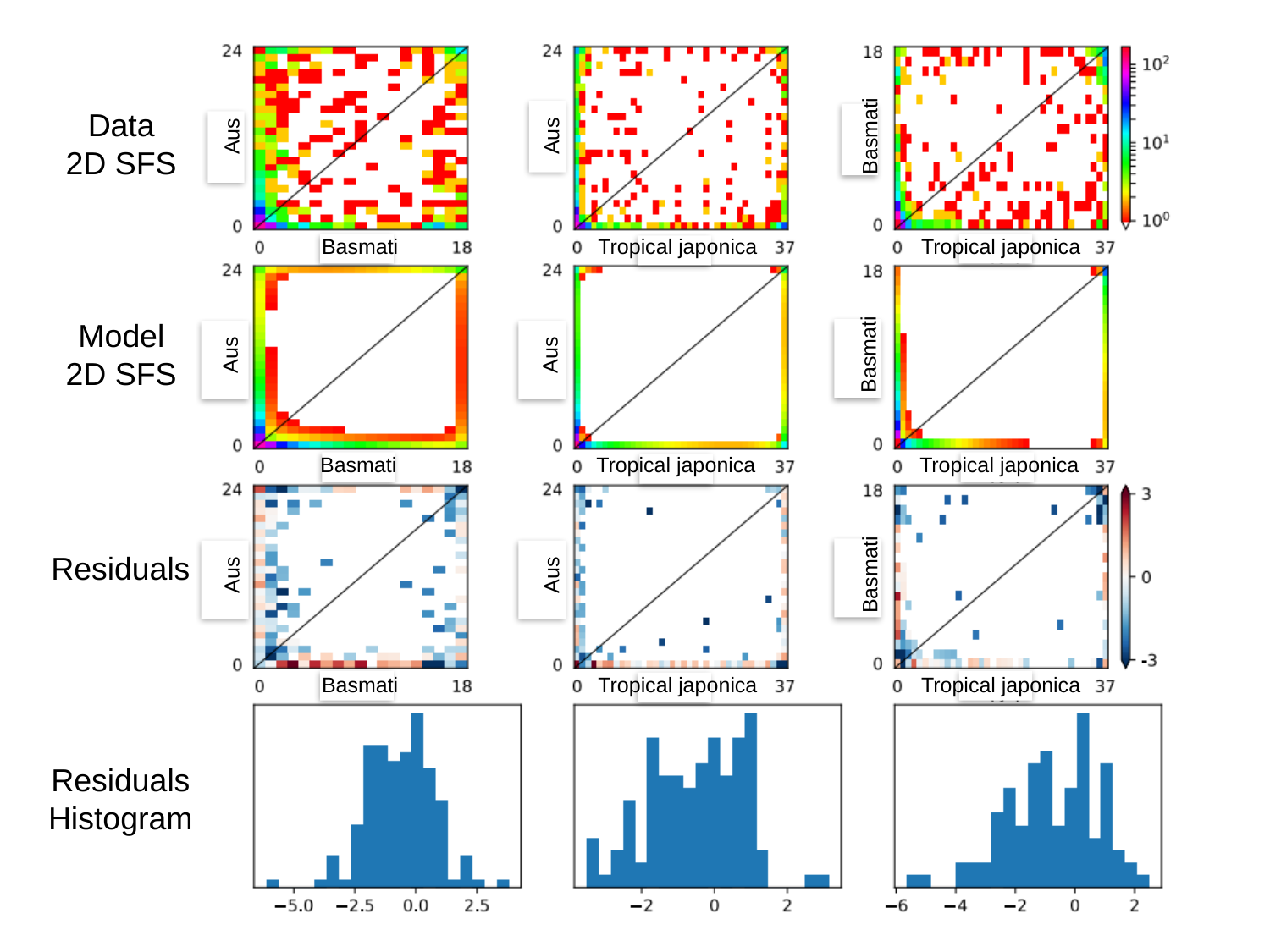

Data
2D SFS
Aus
Aus
Basmati
Tropical japonica
Basmati
Tropical japonica
Model
2D SFS
Aus
Aus
Basmati
Tropical japonica
Basmati
Tropical japonica
Residuals
Aus
Aus
Basmati
Tropical japonica
Basmati
Tropical japonica
Residuals
Histogram
