## Supplementary figures and images for "Nanopore-based genome assembly and the evolutionary genomics of basmati rice"

### Supplemental Figure 1.pptx

#### Slide 1
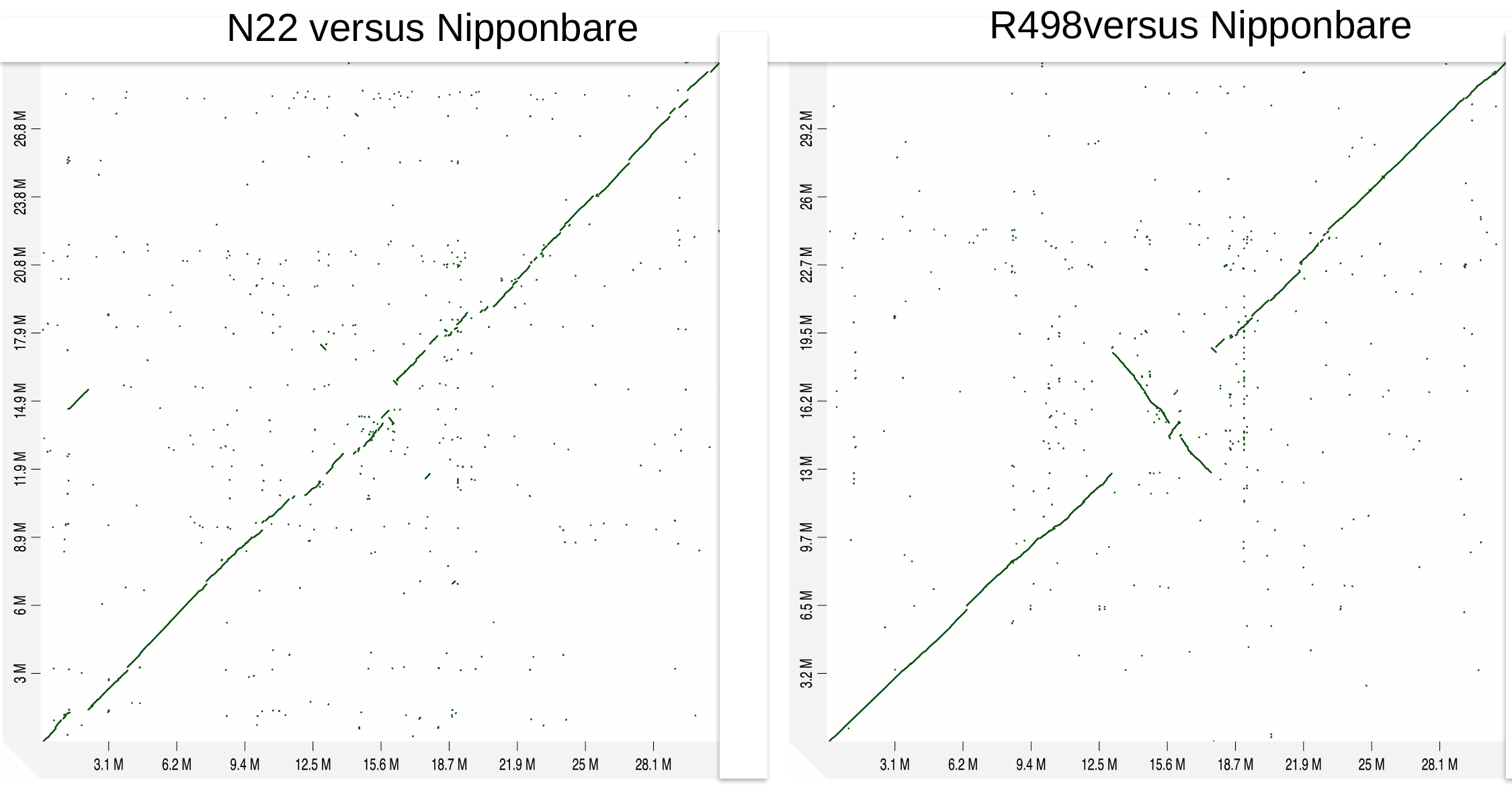

R498versus Nipponbare
N22 versus Nipponbare

### Supplemental Figure 2.pptx

#### Slide 1
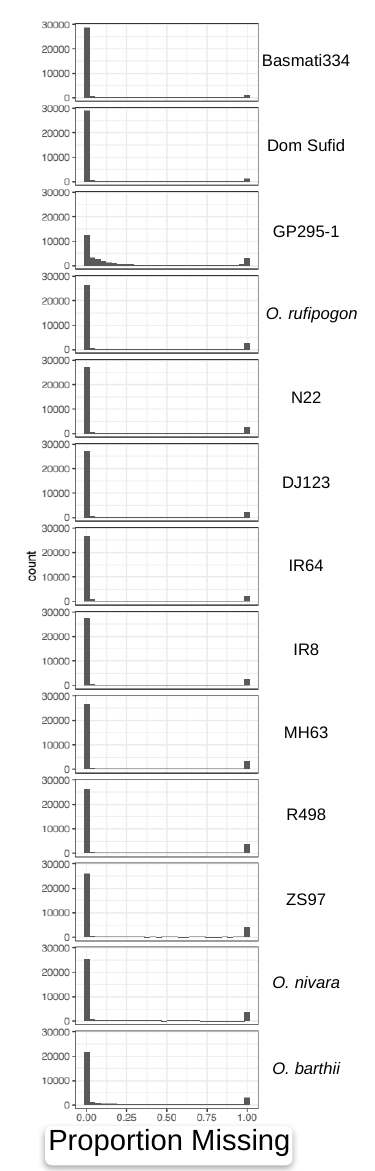

Basmati334
Dom Sufid
GP295-1
O. rufipogon
N22
DJ123
IR64
IR8
MH63
R498
ZS97
O. nivara
O. barthii
Proportion Missing

### Supplemental Figure 3.pptx

#### Slide 1
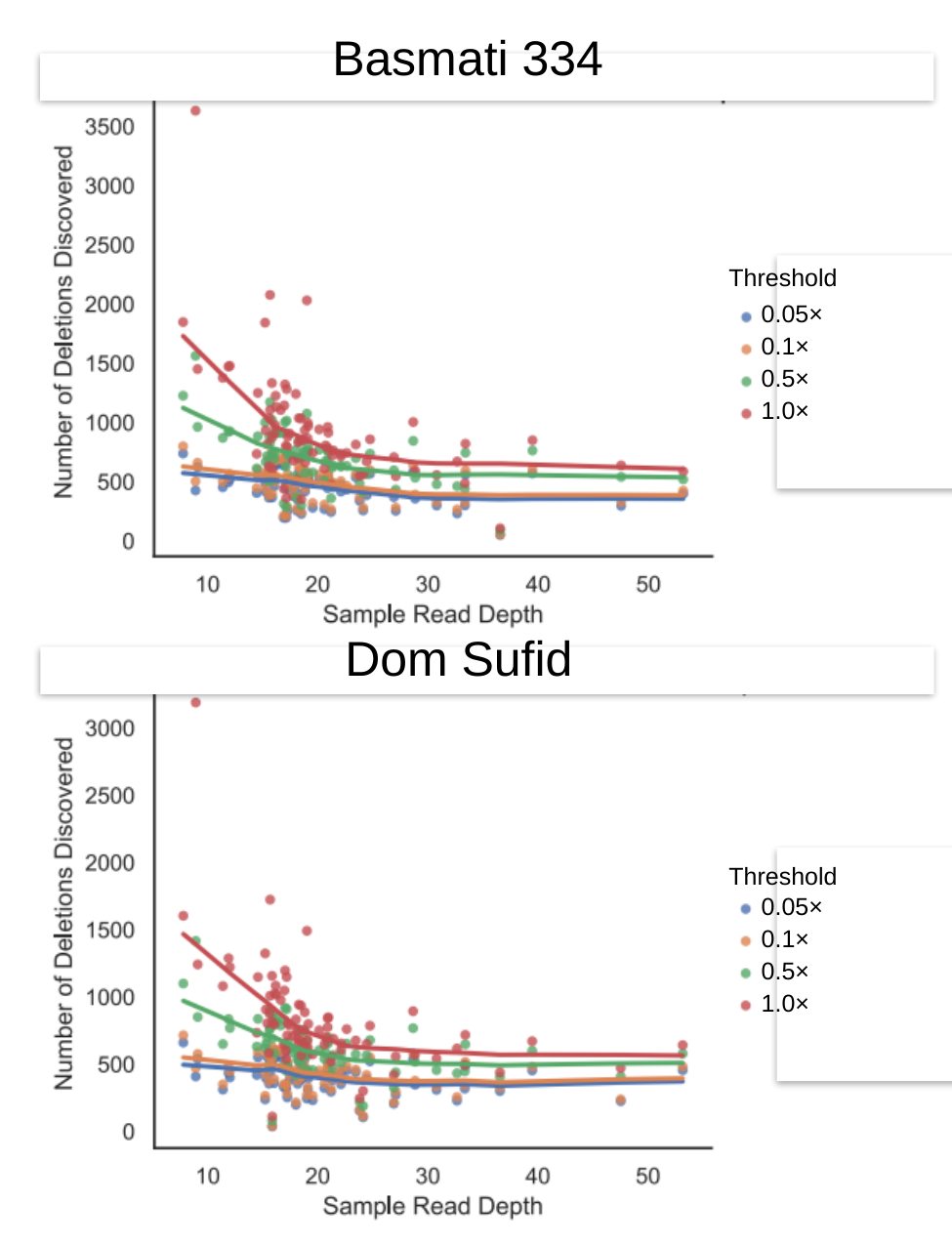

Basmati 334
Threshold
0.05×
0.1×
0.5×
1.0×
Dom Sufid
Threshold
0.05×
0.1×
0.5×
1.0×

### Supplemental Figure 5.pptx

#### Slide 1
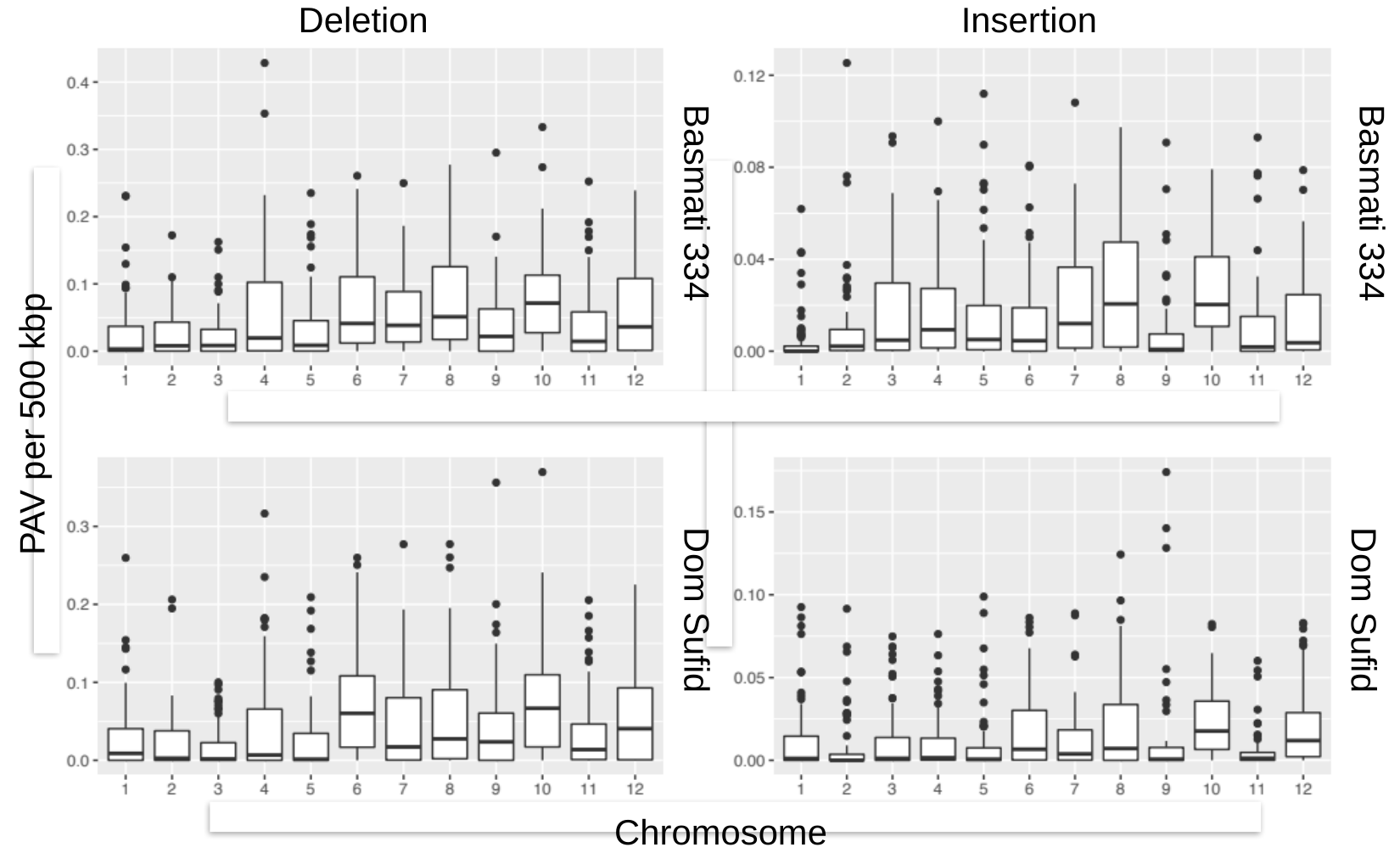

Deletion
Insertion
Basmati 334
Basmati 334
PAV per 500 kbp
Dom Sufid
Dom Sufid
Chromosome

### Supplemental Figure 9.pptx

#### Slide 1
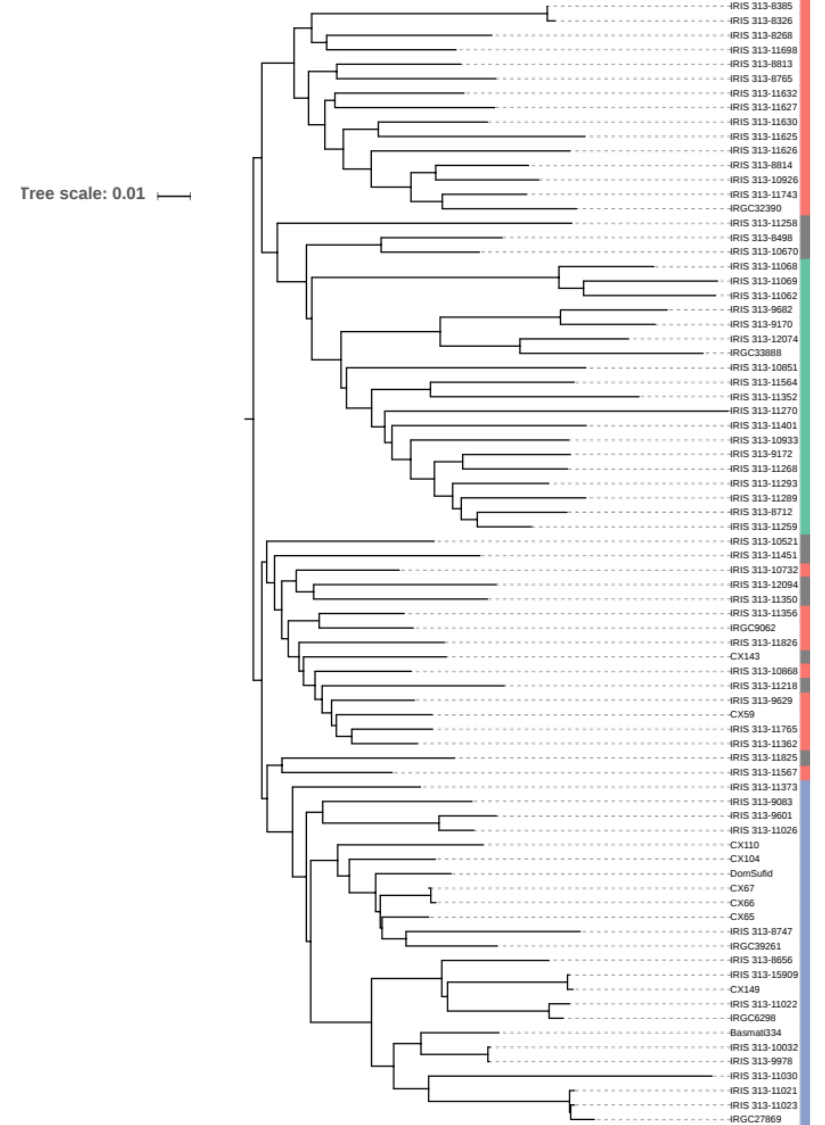
